# Human follicular helper T cell promoter connectomes reveal novel genes and regulatory elements at SLE GWAS loci

**DOI:** 10.1101/2019.12.20.885426

**Authors:** Chun Su, Matthew E. Johnson, Annabel Torres, Rajan M. Thomas, Elisabetta Manduchi, Prabhat Sharma, Carole Le Coz, Michelle E. Leonard, Sumei Lu, Kenyaita M. Hodge, Alessandra Chesi, James Pippin, Neil Romberg, Struan F. A. Grant, Andrew D. Wells

## Abstract

Systemic lupus erythematosus (SLE) is a complex inflammatory disease mediated by autoreactive antibodies that damages multiple tissues in children and adults. Genome-wide association studies (GWAS) have statistically implicated hundreds of loci in the susceptibility to human disease, including SLE, but the majority have failed to identify the causal variants or the effector genes. As a physicochemical approach to detecting functional variants and connecting them to target genes, we generated comprehensive, high-resolution maps of SLE variant accessibility and gene connectivity in the context of the three-dimensional chromosomal architecture of human tonsillar follicular helper T cells (TFH), a cell type required for the production of anti-nuclear antibodies characteristic of SLE. These spatial epigenomic maps identified a shortlist of over 400 potentially functional variants across 48 GWAS-implicated SLE loci. Twenty percent of these variants were located in open promoters of highly-expressed TFH genes, while 80% reside in non-promoter genomic regions that are connected in 3D to genes that likewise tend to be highly expressed in TFH. Importantly, we find that 90% of SLE-associated variants exhibit spatial proximity to genes that are not nearby in the 1D sequence of the genome, and over 60% of variants ‘skip’ the nearest gene to physically interact only with the promoters of distant genes. Gene ontology confirmed that genes in spatial proximity to SLE variants reside in highly SLE-relevant networks, including accessible variants that loop 200-1000 kb to interact with the promoters of the canonical TFH genes *BCL6* and *CXCR5*. CRISPR-Cas9 genome editing confirmed that these variants reside in novel, distal regulatory elements required for normal *BCL6* and *CXCR5* expression by T cells. Furthermore, SLE-associated SNP-promoter interactomes implicated a set of novel genes with no known role in TFH or SLE disease biology, including the homeobox-interacting protein kinase HIPK1 and the Ste kinase homolog MINK1. Targeting these kinases in primary human TFH cells inhibited production of IL-21, a requisite cytokine for production of class-switched antibodies by B cells. This 3D-variant-to-gene mapping approach gives mechanistic insight into the SLE-associated regulatory architecture of the human genome.

## INTRODUCTION

GWAS has been an important tool in understanding the genetic basis of complex, heritable diseases and traits. However, GWAS is typically powered to identify relatively large blocks of the genome containing dozens to hundreds of single nucleotide polymorphisms (SNP) in linkage disequilibrium (LD), any one of which could be responsible for the association of the entire locus with disease susceptibility. Moreover, ∼90% of GWAS-implicated SNP are intergenic or intronic, and do not affect the coding sequence of proteins. Therefore, the location of the GWAS signal *per se* does not identify the culprit gene(s). Examples of this are the *FTO* GWAS signal in obesity^1, 2^, and the *TCF7L2* GWAS signal in type 2 diabetes^3^, in which each suspected causal variant resides in an intron of the local gene, but were shown instead to regulate expression of the distant genes *IRX3/5* and *ACSL5*, respectively.

Systemic lupus erythematosus (SLE) is a complex inflammatory disease mediated by autoreactive antibodies that damage skin, joints, kidneys, brain and other tissues in children and adults^4^. An important inflammatory leukocyte required for the development of SLE is the follicular helper T cell (TFH). TFH differentiate from naïve CD4+ T cells in the lymph nodes, spleen, and tonsil, where they license B cells to produce high affinity protective or pathogenic antibodies^5, 6^. Given the central role for TFH in the regulation of humoral immune responses, genetic susceptibility to SLE is highly likely to manifest functionally in this immune cell population.

GWAS has associated over 60 loci with SLE susceptibility to date^7, 8^, but this represents thousands of SNP in LD of which any could potentially contribute to disease. Given the paucity of immune cell eQTL data represented in GTEx, we mapped the open chromatin landscape of TFH cells from human tonsil to identify potentially functional SLE variants, and conducted a genome-wide, promoter-focused Capture-C analysis of chromatin contacts at nearly all of the ∼41,000 annotated human protein-coding and non-coding genes at ∼270 bp resolution to map these variants to the genes they likely regulate in this disease-relevant cell type. This approach leverages the power of existing SLE genetic knowledge, using the location of common variants that have already been strongly associated with SLE pathogenesis to identify (*via* assessment of linkage disequilibrium) putative disease-associated regulatory elements. We then utilize high-resolution, promoter-focused Capture-C to physically connect these variants to their target genes, in line with our previous, comparable approach to studying bone mineral density loci^14^. This approach only requires three replicate samples to make valid interaction calls, and it does not require material from SLE patients or genotyped individuals. By design, we are not studying the effect of a variant in the system, but rather, we are using reported variants as ‘signposts’ to identify potential enhancers in healthy tissue. The variant connectomes in turn lead us to putative effector genes that warrant further follow up. This study shows that the majority of accessible SLE-associated variants do not interact with the nearest promoter, but are instead connected to more distant genes, many of which have known roles in T cell and TFH biology. Using CRISPR/CAS9 genome editing, we go on to validate several of these SLE-associated open chromatin regions, revealing a requisite role in regulating their connected genes. Finally, we experimentally verified that two novel kinases implicated by this variant-to-gene mapping approach are required for TFH differentiation and/or function, and therefore represent potential novel drug targets for SLE and other antibody-mediated diseases.

## RESULTS

### Comparative open chromatin landscapes of naïve CD4+ T cells and TFH from human tonsil

The vast majority (>90%) of the human genome is packed tightly into cellular chromatin and is not accessible to the nuclear machinery that regulates gene expression^9^. Consequently, >95% of transcription factor and RNA polymerase occupancy is concentrated at regions of open chromatin^9^, and thus a map of accessible chromatin in a given cell type essentially defines its potential gene regulatory landscape. As a step toward defining the disease-associated regulatory architecture of the complex heritable autoimmune disease SLE, we focused on human follicular helper CD4+ T cells (TFH), which are required for the production of pathogenic antibodies by autoreactive B cells during the development of SLE^4^. Tonsillar TFH are derived from naïve CD4+ T cell precursors, and represent a population of cells in healthy subjects that are actively in the process of helping B cells to produce high-affinity, class-switched antibodies. We sorted naive CD4+CD45RO-T cells and differentiated CD4+CD45RO+CD25-CXCR5^hi^PD1^hi^ TFH^10^ from human tonsil and generated open chromatin maps of both cell types from three donors using ATAC-seq^11^. A binary peak calling approach identified a total of 91,222 open chromatin regions (OCR), 75,268 OCR in naïve CD4+ cells and 74,627 OCR in TFH cells (**Supplemental Table 1**). Further quantitative analysis of the accessibility signal at these OCR revealed a similar overall degree of genomic accessibility (∼1.4%) in both cell types (**Supplemental Figure 1**). However, the differentiation of naïve CD4+ T cells into TFH is associated with remodeling of 22% of the T cell open chromatin landscape, with 11,228 OCR becoming more accessible, and 8,804 OCR becoming less accessible, in the TFH lineage (**Figure 1A, Supplemental Table 1**). Among all 20,032 differentially accessible regions, 20.5% (4100) reside in promoters, and the genes driven by these differentially accessible promoters tended to be differentially expressed between TFH and naïve CD4+ cells as assessed by microarray (**Figure 1B**, GSEA enrichment p<0.05, absNES>3.5, **Supplemental Table 2**). The functions of genes that are remodeled and upregulated in TFH were significantly enriched for CD28 costimulatory, G-protein, Rho GTPase, semaphorin, and TLR signaling pathways (hypergeometric test, FDR<0.05, **Supplemental Figure 2A**). Conversely, gene promoters that become less accessible upon TFH differentiation are enriched for pathways involved in chemokine and G protein-coupled signaling (**Supplemental Figure 2B**). These data show that global chromatin remodeling dynamics faithfully reflect dynamic changes in gene expression during the differentiation of follicular helper T cells from their naïve precursors.

**FIGURE 1.**
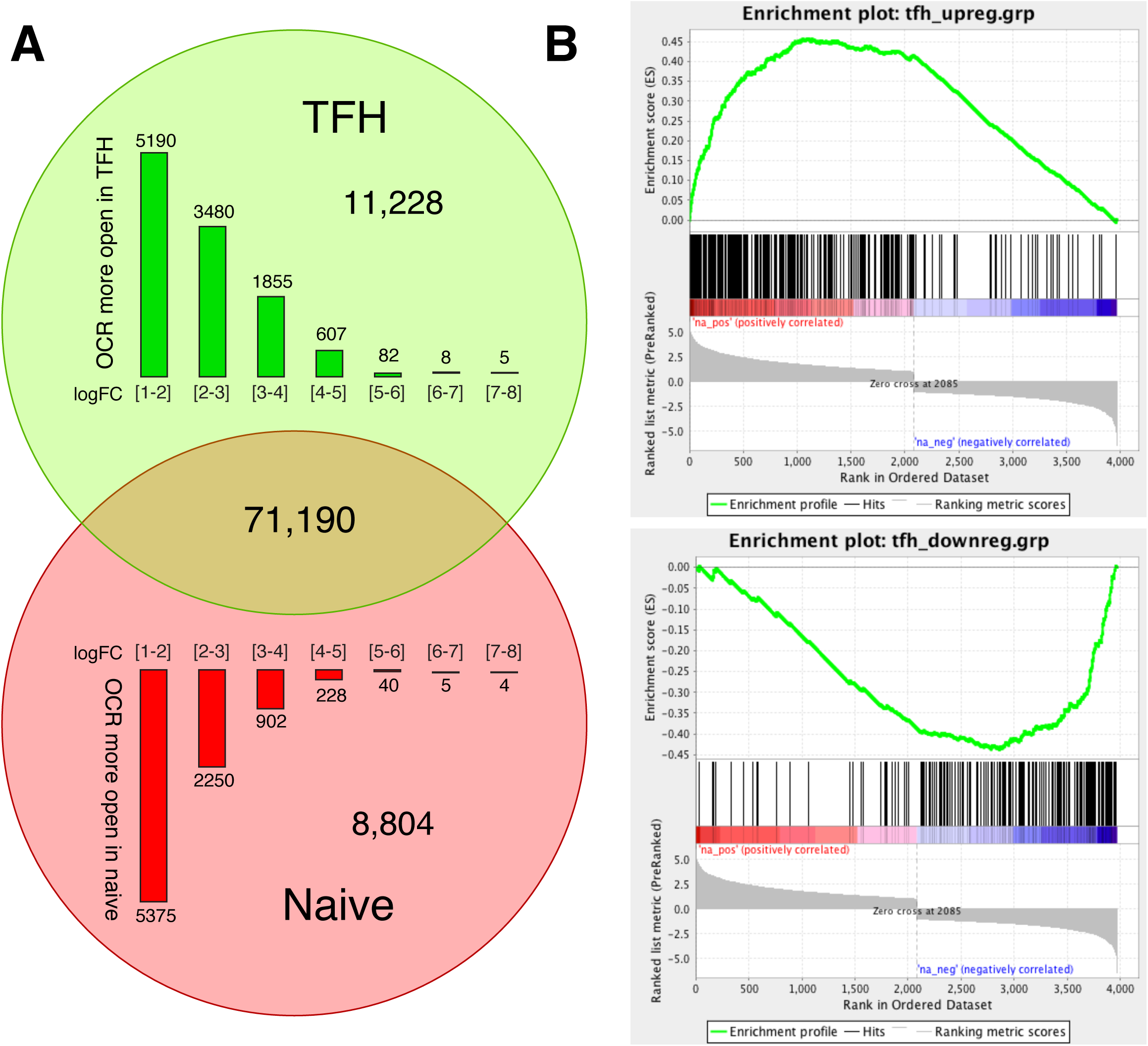
ATAC-seq analysis of open chromatin landscapes in naïve and follicular helper T cells from human tonsil. a. Quantitative differences between naive and follicular helper T cell open chromatin landscapes. A total of 91,222 OCR were used as reference for differential analysis of genome accessibility. The number of statistically up- (green) or down- (red) regulated OCR in TFH compared to naïve is shown as a Venn diagram and also plotted as a function of log2 fold change. b. GSEA enrichment analysis of genes with differential promoter accessibility at promoter regions. The log2 fold change in expression between naïve and follicular helper T cells was used to generate the pre-ranked list for GSEA.

### Open chromatin mapping of disease-relevant tissue implicates causal disease variants

Of the 7662 sentinel and proxy SNPs currently implicated by GWAS in SLE (r^2^>0.4)^7, 8^, 432 (5.6%) reside in 246 open chromatin regions in either naïve CD4+ T cells or TFH (**Supplemental Table 3**). Of these, 345 SNPs (80%) are in open chromatin shared by both cell types, 39 are in naive-specific OCR, and 48 reside uniquely within the TFH open chromatin landscape (**Figure 2A**). Altogether, 91% (393 of 432) of the accessible SLE SNPs identified in this study reside in the open chromatin landscape of TFH cells. To explore the potential significance of these open chromatin-implicated variants, we first focused on the 132 SLE proxy SNPs that reside in open promoters of protein-coding genes in TFH. Of the 64 genes containing one or more open promoter variants, 62 are expressed in TFH as assessed by microarray (**Supplemental Table 2**). Moreover, the set of genes with accessible SLE variants in their promoters are expressed more highly in TFH than the set of all genes, or compared to a random sample of genes with open promoters in TFH (**Figure 2B**). Eighty-three SLE proxy SNPs reside in promoters of 36 genes in the top 75% expression quantile, and 43 SLE proxy SNPs reside in the promoters of 18 genes in the 50-75% expression quantile. Thus, nearly 93% (123 of 132) of open promoter SLE proxy variants are positioned at genes moderately to highly expressed in TFH. Ingenuity pathway analysis (IPA) found that these TFH expressed genes are enriched for factors involved in systemic lupus erythematosus (*P*=4.6×10^-9^), systemic autoimmunity (p=6.0×10^-8^), and rheumatic disease (*P*=2.7×10^-7^) (**Figure 2C**), including *PTPRC* (CD45), *TCF7* (TCF1), *IRF5, IFNLR1, TYK2, ELF1, IKZF2* and *JAK2*. This set of highly-expressed TFH genes with SLE variants in their promoters also includes *DHCR7* and *NADSYN1*, enzymes involved in biogenesis of vitamin D, a process known to play an important role in autoimmune disease susceptibility^12^. These results indicate that open chromatin landscapes in disease-relevant cell types represent a highly specific filter through which putatively functional SLE variants can be distinguished from the thousands of proxy SNPs to the sentinels implicated by GWAS.

**FIGURE 2.**
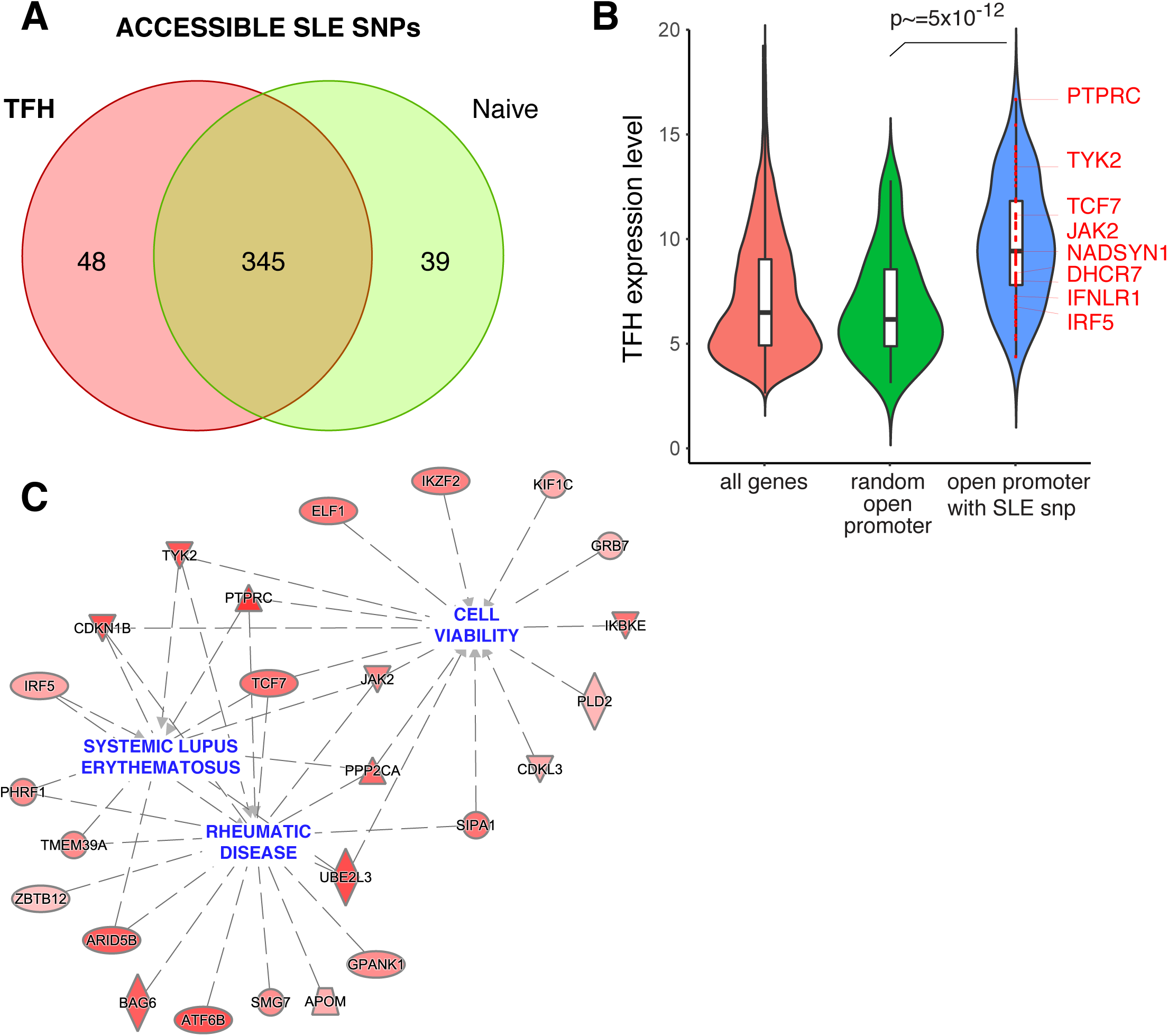
Genes harboring accessible SLE variants in naïve and follicular helper T cells. a. Comparison of accessible SLE SNPs between TFH and naïve tonsillar T cells. b. Comparison of the expression levels of genes with accessible SLE SNPs in their promoters in TFH vs. all genes or a random sample of genes with no accessible SLE SNPs in their promoters. A Wilcoxon rank test was performed to evaluate the significance of differential expression between gene sets. c. Ingenuity disease network for the genes with accessible SLE variants at promoters. The color gradient represents the log2 fold change IN expression in TFH compared to naïve T cells.

### Analysis of the three-dimensional promoter connectome structure in TFH cells

It is relatively clear how genetic variation at a promoter might influence expression of the downstream gene. However, ∼70% of the accessible SLE SNPs in TFH cells are intronic, intergenic, or otherwise not in proximity to a promoter, so how these variants regulate the expression of specific TFH genes is not clear from 1-dimensional open chromatin mapping alone. To explore the role that accessible, non-coding SLE-associated variants play in the disease-related regulatory architecture of the human genome, we derived genome-wide, three-dimensional promoter contact maps of naive CD4+ T cells and differentiated follicular helper CD4+ T cells from human tonsil using promoter-focused Capture-C technology. Our chromosome conformation capture (3C)-based approach is a high-resolution, large-scale modification of Capture-C^13^ that involves massively parallel, hybridization-based enrichment of 41,970 targeted promoters associated with 123,526 currently annotated transcripts (gencode v19) covering 89% of protein-coding mRNA genes and 59% of non-coding (anti-sense RNA, snRNA, miRNA, snoRNA and lincRNA) genes in the human genome^14^. As in standard capture-C approaches, valid hybrid reads derived from ligation of distant fragments with bait fragments were preprocessed using HiCUP^15^, and significant promoter-interacting regions (PIR, score >5) were called using CHiCAGO^16^. Unlike promoter capture Hi-C^17^, our method employs the 4-cutter DpnII to generate 3C libraries with a 270 bp median resolution, ∼9-fold higher than the 2300 bp median resolution of the HindIII-based 3C libraries generated in HiC and capture-HiC approaches. This resolution allows mapping of interactions between promoters and distal regulatory elements to within a span of two nucleosomes. This precision comes at the expense of power, in that sequencing reads are distributed across more fragments, leaving fewer reads available per fragment to call significant promoter interactions. To circumvent this problem, we called promoter interactions both at high resolution (single-fragment) and at lower resolution (four-fragment) after an *in silico* fragment concatenation step. Combination of both sets of calls allows this method to benefit from the precision of single DpnII fragment analysis and the power of lower resolution analyses at farther distances to assemble comprehensive, 3D promoter contact maps for the human genome (**Figure 3A**).

**FIGURE 3.**
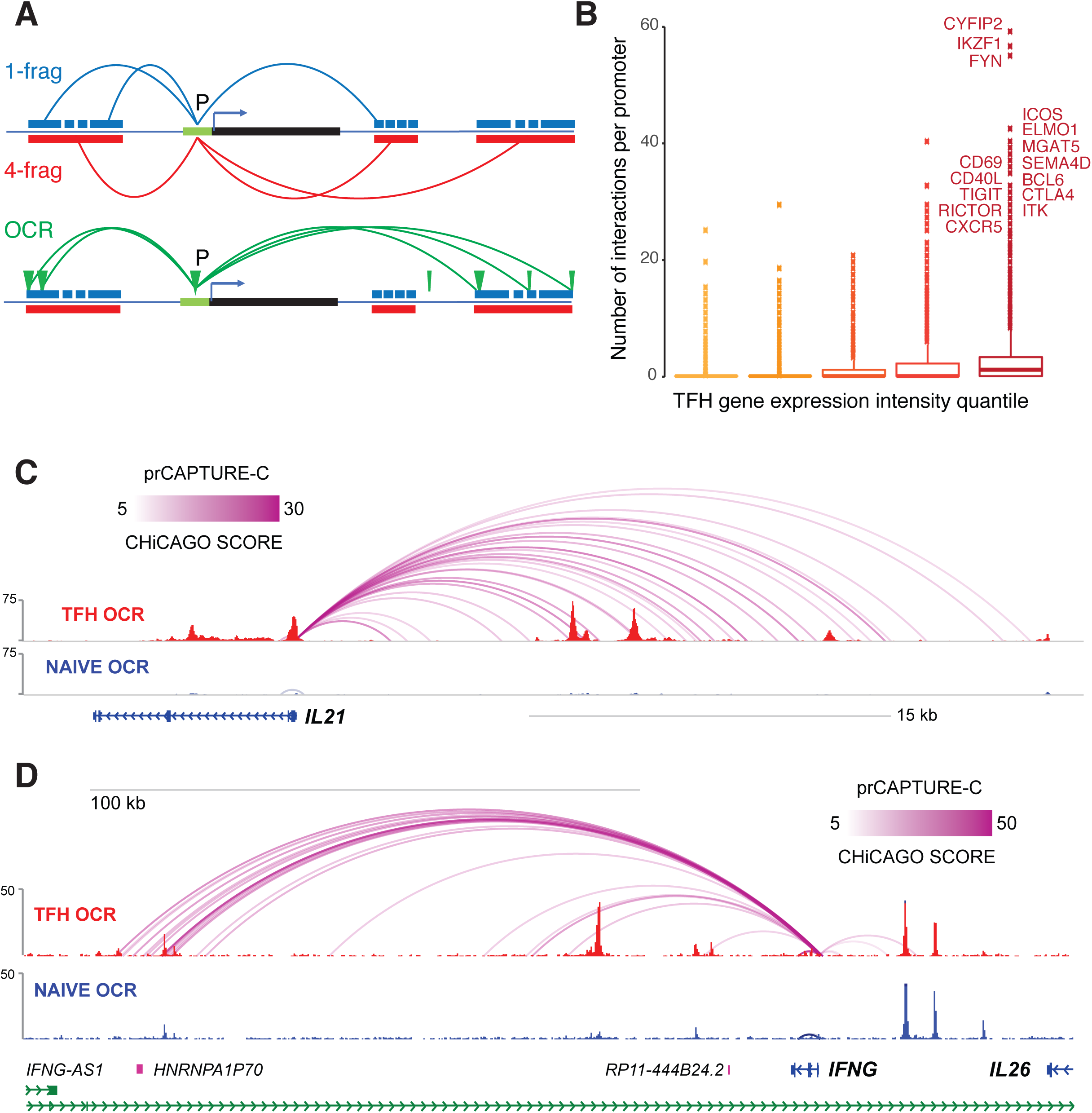
High-resolution, fragment-based Capture-C analysis of promoter connectomes in naïve and follicular helper T cells. a. Cartoon depicting the approaches for 1 DpnII fragment promoter interaction analysis, 4 DpnII fragment promoter interaction analysis, and promoter-OCR interaction analysis. b. The relationship between the number of interactions per gene promoter and expression of the corresponding gene is shown. Gene expression was binned into the lowest 20th, 20-40th, 40-60th, 60-80th and >80th percentiles. Lower and upper boxplot hinges correspond to the first and third quartiles, and outliers were defined as > 1.5 * IQR from the hinge. WashU browser depictions of fragment-based promoter interactions and ATAC-seq accessibility at the IL21 (c) and IFNG (d) loci in TFH (red) and naive T cells (blue). Color gradients represent the CHiCAGO scores.

We detected a similar number of significant promoter interactions (CHiCAGO score >5) in both cell types - 255,238 in naive CD4+ T cells and 224,263 in TFH - with the vast majority (>99%) being intra-chromosomal (*in cis*). About 20% of total interactions were between two promoters, while 80% of interactions were between a promoter and an intergenic or intronic region (**Supplemental Table 4**). Ninety percent of promoters were found engaged in at least one stable interaction with another genomic region. Of these promoters, over 80% were connected to only one distal genomic region, indicating that most promoters in these cell types exhibit very low spatial complexity. However, ∼1% of all promoters exhibited significant spatial complexity, interacting with four or more distal regions, with some promoters engaging in as many as 70 interactions with distal regions. The number of connections per promoter correlated with the level of gene expression in both cell types, with the most interactive promoters belonging to highly-expressed genes with known roles in TFH function (**Figure 3B**). Two examples are the *IL21* and *IFNG* promoters, which are expressed and show complex connectomes in TFH but not naïve cells (**Figures 3C** and **D**). Promoter-interacting regions in both cell types were enriched 3-fold for open chromatin, and 2-fold for chromatin signatures associated with active transcription, such as H3K27ac, H3K4me1, H3K4me3 (**Figure 4A** and **Supplemental Figure 3A**). Conversely, PIR in both cell types were depleted of the silencing marks H3K27me3 and H3K9me3 (**Figure 4A** and **Supplement Figure 3A**). Together, these trends indicate that the promoter contacts captured by this approach preferentially represent the active regulatory architectures of the associated genes.

**FIGURE 4.**
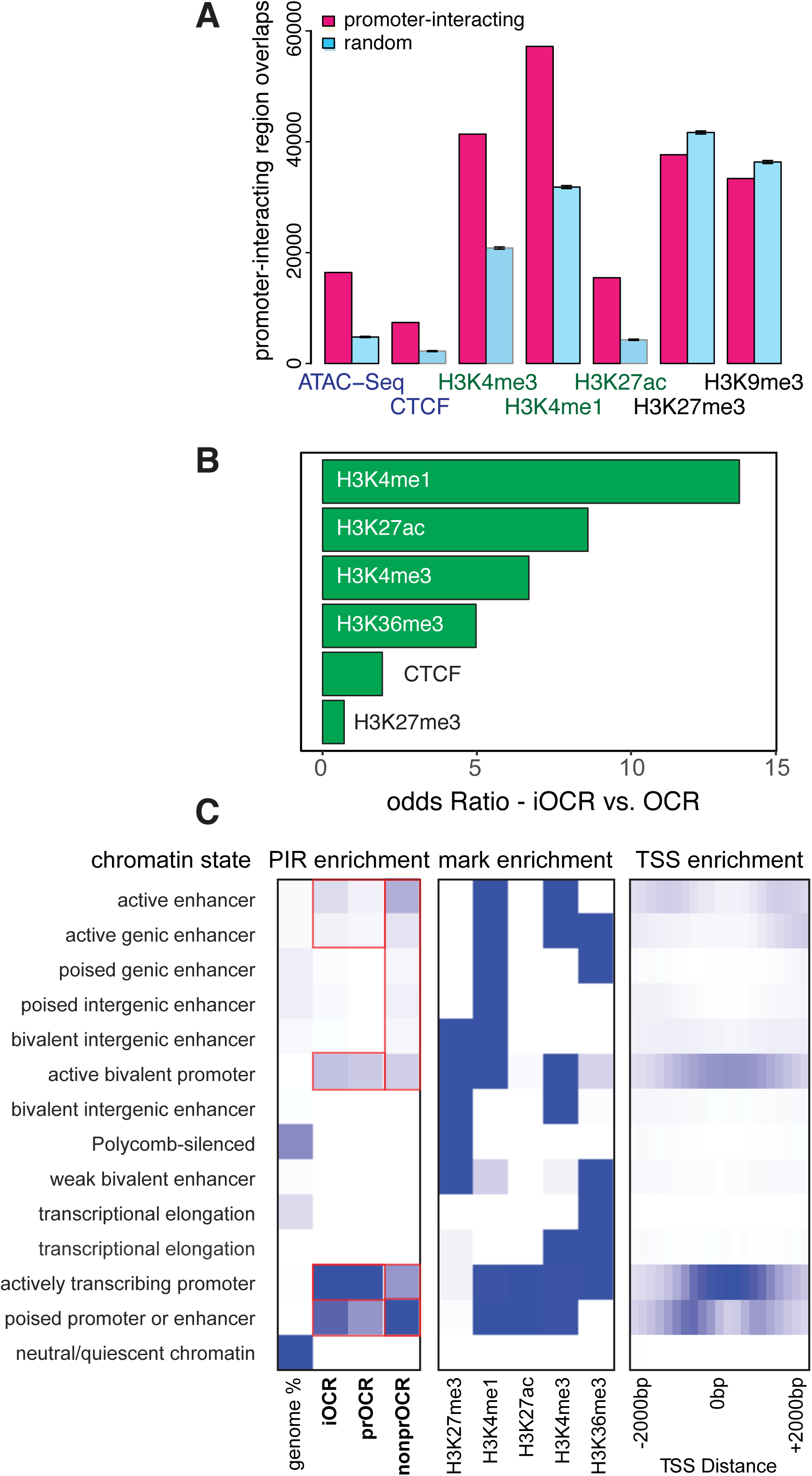
Enrichment of chromatin signatures at promoter interacting regions. a. PIR enrichment for genomic features compared with distance-matched random regions in naive T cells. Error bars show SD across 100 draws of non-significant interactions. b. Feature enrichment at promoter-interacting OCR (iOCR) compared to a random sample of non-promoter-interacting OCR in naive T cells. c. Enrichment of iOCR within chromHMM-defined chromatin states and TSS neighborhood in naive T cells. ChromHMM 15-state models defined on the basis of 5 histone modifications (H3K4me1, H3K4me3, H3K27me3, H3K27ac and H3K36me3) are shown in the middle panel, with blue color intensity representing the probability of observing the mark in each state. The heatmap to the left of the emission parameters displays the overlap fold enrichment for iOCR in promoters (prOCR) and non-promoter iOCR (nonprOCR), while the heatmap to the right shows the fold enrichment for each state within 2 kb around a set of TSS. Blue color intensity represents fold enrichment.

### The promoter-open chromatin connectome of TFH cells

To further explore the regulatory nature of the spatial connections between promoters and other genomic regions in the nucleus, we focused on contacts between promoters and open chromatin regions, as the biochemical processes that regulate transcription occur largely at accessible DNA^9^. Instead of using standard fragment-based interactions^13^, we used a feature-based calling approach to define interactions between promoters (−1500 to +500 from a TSS) and OCR, combining calls at both one- and four-fragment resolution to generate genome-wide, open chromatin-promoter interaction landscapes in naïve and follicular helper CD4+ T cells (**Figure 3A**). In total, we detected 71,137 *cis*-interactions between accessible promoters and open chromatin among both cell types, involving 34% of the total open chromatin landscape identified by our ATAC-seq analyses (**Supplemental Table 5**). We define these 31,404 promoter-interacting OCR as iOCR. Roughly half of iOCR (15,109, 48%) are located in intergenic or intronic regions relatively far from genes, while the other 16,295 promoter-interacting OCR (52%) are located in the promoters of other distant genes. The distance between promoter and promoter-interacting OCR pairs ranged from a few hundred base pairs to over a megabase, with a median of ∼112 kb for both categories. Remarkably, while OCR in general are enriched for active chromatin marks^9, 11^, we find that iOCR are even more highly enriched for enhancer signatures compared to OCR that are not contacting a promoter (14-fold for H3K4me1, 9-fold for H3K27ac, 7-fold for H3K4me3, Fisher test p<2×10^-16^, **Figure 4B** and **Supplemental Figure 3B**. Chromatin state modeling (chromoHMM^18^) revealed that all iOCR are enriched at active promoters, bivalent promoters, and active enhancers, as defined by histone modification ChIP-seq in both naïve and TFH cells^19–21^ (**Figure 4C** and **Supplemental Figure 3C**). Open promoter regions involved in promoter interactions (prOCR) were more specifically enriched with active promoter signatures, while promoter-interacting OCR located in intergenic/intronic space (nonprOCR) were more specifically enriched at poised and active enhancers (**Figure 4C**). These results indicate that promoter-connected OCR are biochemically distinct from OCR not connected to a promoter, and that this promoter-Capture-C approach enriches for genomic elements that are actively engaged in gene regulation.

Using this open chromatin-promoter interaction landscape, we were able to connect the promoters of 18,669 genes (associated with 79,330 transcripts) to their corresponding putative regulatory elements, representing 145,568 distinct gene-OCR interactions. Roughly half (47%, 68,229) of these interactions occur in both naive and TFH cells, while 24% (34,928) occur uniquely in naïve cells, and 29% (42,411) are only found in TFH (**Figure 5A**). Overall, 91% of OCR-connected genes (17,021) exhibit at least one differential promoter-OCR interaction in naïve vs. follicular helper T cells. The majority (82%) of OCR-connected genes were incorporated into regulatory structures consisting of more than one distal regulatory region in naïve and follicular helper T cells. On average, each of these connected genes interact with 6 OCR in both naïve CD4+ T cells and follicular helper T cells (4 in median, **Supplemental Figure 4**), with 10% of these genes involved in 13 or more interactions with distal OCR. More interestingly, the degree of spatial connectivity exhibited by a promoter tends to positively correlate with the level of gene expression in a lineage-specific manner (**Figure 5B** and **C**). The common, highly-connected promoters in both cell types drive the expression of genes involved in cell cycle, DNA organization and repair, protein and RNA biogenesis and trafficking, and TCR signaling (**Supplemental Figure 5**). In addition, highly interactive promoters in naïve cells are involved in quiescence, signal transduction and immune function (*e.g*., *FOXP1, CCR7, IKZF1, CD3, FYN, GRB2, GRAP2, BIRC2/3;* **Figure 5B**), while gene promoters that exhibit complex regulatory architectures in TFH are highly expressed in TFH and are involved in TFH and T cell differentiation, survival, homing, and function (*e.g*., *BCL6, CXCR5, CD40L, CTLA4, ICOS, CD2, CD3, CD28, CD69, TCF7, NFAT1, BATF, ITK, IKZF2, IKZF3, IL21R, FAS*; **Figure 5C**). An example is the *CD28-CTLA4-ICOS* multi-locus region. In naïve CD4+ T cells, which express *CD28* but not *CTLA4* or *ICOS*, the *CD28* promoter is engaged in multiple interactions with 8 downstream regions of open chromatin (**Figure 5D**, blue), while the *CTLA4* and *ICOS* promoters are much less interactive. In TFH, which express all three genes, the *CD28*, *CTLA4*, and *ICOS* promoters adopt extensive, *de novo* spatial conformations involving contacts with more than two dozen putative regulatory elements within the TFH-specific open chromatin landscape (**Figure 5D**, red). Together, these results reveal major restructuring of the T cell gene regulatory architecture that occurs as naïve helper T cells differentiate into follicular helper T cells, and indicate that complex, three-dimensional regulatory architectures are a feature of highly expressed, lineage-specific genes involved in specialized immune functions in this disease-relevant cell types.

**FIGURE 5.**
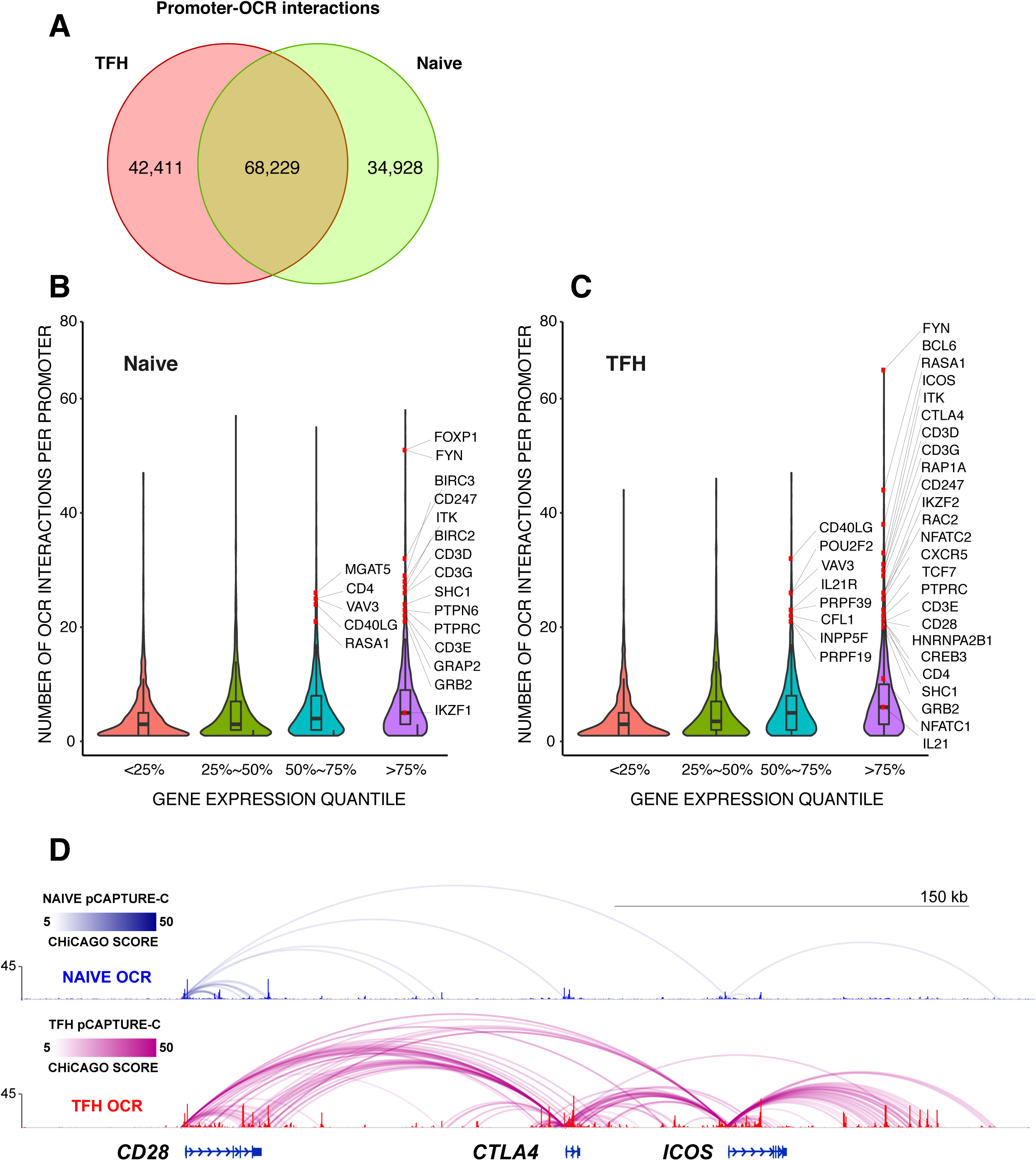
Analysis of promoter-open chromatin connectomes in naïve and follicular helper T cells. a. Comparison of promoter-OCR interactions between TFH and naïve T cells. Relationship between the number of promoter-OCR interactions at a gene and its corresponding expression level in naive T cells (b) and TFH (c) are depicted with genes with recognized functions in naive and TFH labeled. d. Promoter-OCR interactions and ATAC-seq accessibility at the CD28, CTLA4 and ICOS loci in TFH (red) and naive T cells (blue).

### Disease-associated variant-to-gene mapping for SLE

The open chromatin landscape of follicular helper T cells from the three healthy tonsils studied contained 393 accessible genomic regions that harbor SLE disease variants (**Supplemental Table 3**), representing the TFH component of the potential *cis*-regulatory landscape of SLE. While 33% of accessible variants (132 proxy SNPs) reside in promoters, 67% of accessible SLE SNPs (N=261) are in non-promoter regions. Therefore, the role these non-promoter variants play in gene transcription, and which genes they control, is not clear from these one-dimensional epigenomic data. To determine whether spatial proximity of a gene to an open SLE SNP in three dimensions is a predictor of its role in TFH and/or SLE, we explored the 3D *cis*-regulatory architecture of SLE genetic susceptibility based upon open chromatin region interaction landscape generated in TFH cells, effectively mapping 256 open SLE variants (69 sentinels, r^2^>0.4) to 330 potential target genes (1107 SNP-target gene pairs). This 3D variant-to-gene map shows that only ∼9% (22) of the SLE variants that reside in TFH open chromatin interact exclusively with the nearest gene promoter (**Supplemental Table 6**). An example of this category is rs35593987, a proxy to the SLE sentinel SNP rs11889341 and rs4274624 that resides in a TFH OCR and loops ∼99 kb to interact with the *STAT4* promoter (**Figure 6A**). Another ∼30% (75) of open SLE variants interact with nearest promoter, but also with the promoters of more distant genes (**Figure 6A, Supplemental Table 6**). An example of this category is rs112677036, a proxy to the SLE sentinel SNP rs12938617 that resides in the first intron of *IKZF3*, interacts with nearby *IKZF3* promoter, but also interacts with promoters of two 157kb upstream genes *PGAP3* and *ERBB2* (**Figure 6B**). Remarkably, over 60% of all open SLE variants (159) ‘skip’ the nearest gene to interact with at least one distant promoter (**Supplemental Table 6**). Examples of this most abundant category are rs34631447, a proxy to the SLE sentinel rs6762714 SNP that resides in open chromatin in the sixth intron of the *LPP* locus, and the intergenic rs527619 and rs71041848 SNPs proxy to SLE sentinel rs4639966. Our 3D regulatory map in TFH cells demonstrates that the ‘*LPP’* variant in fact does not interact with the *LPP* promoter, but instead is incorporated into a chromosomal loop structure spanning over 1 Mb that positions it in direct, spatial proximity to the promoter of *BCL6*, the ‘master’ transcription factor of follicular helper T cells^22–26^ (**Figure 6C**). Similarly, the OCR containing the SLE proxies rs527619 and rs71041848 does not contact the nearby *TREH* gene, but instead loops to interact with the promoter of the TFH-specific chemokine receptor gene *CXCR5*, nearly 200 kb away (**Figure 6D**). Other relevant examples of this class of SLE SNPs are rs3117582 and rs7769961, proxies to SLE sentinel SNP rs1150757 and rs9462027, respectively. These SNPs in TFH open chromatin were found in contact with *LSM2* and *SNRPC* (**Supplemental Figure 6**), 35 to 150 kb away. Both of these genes encode proteins that participate in the processing of nuclear precursor messenger RNA splicing, and are frequently the targets of autoantibodies produced by patients with SLE^27, 28^.

**FIGURE 6.**
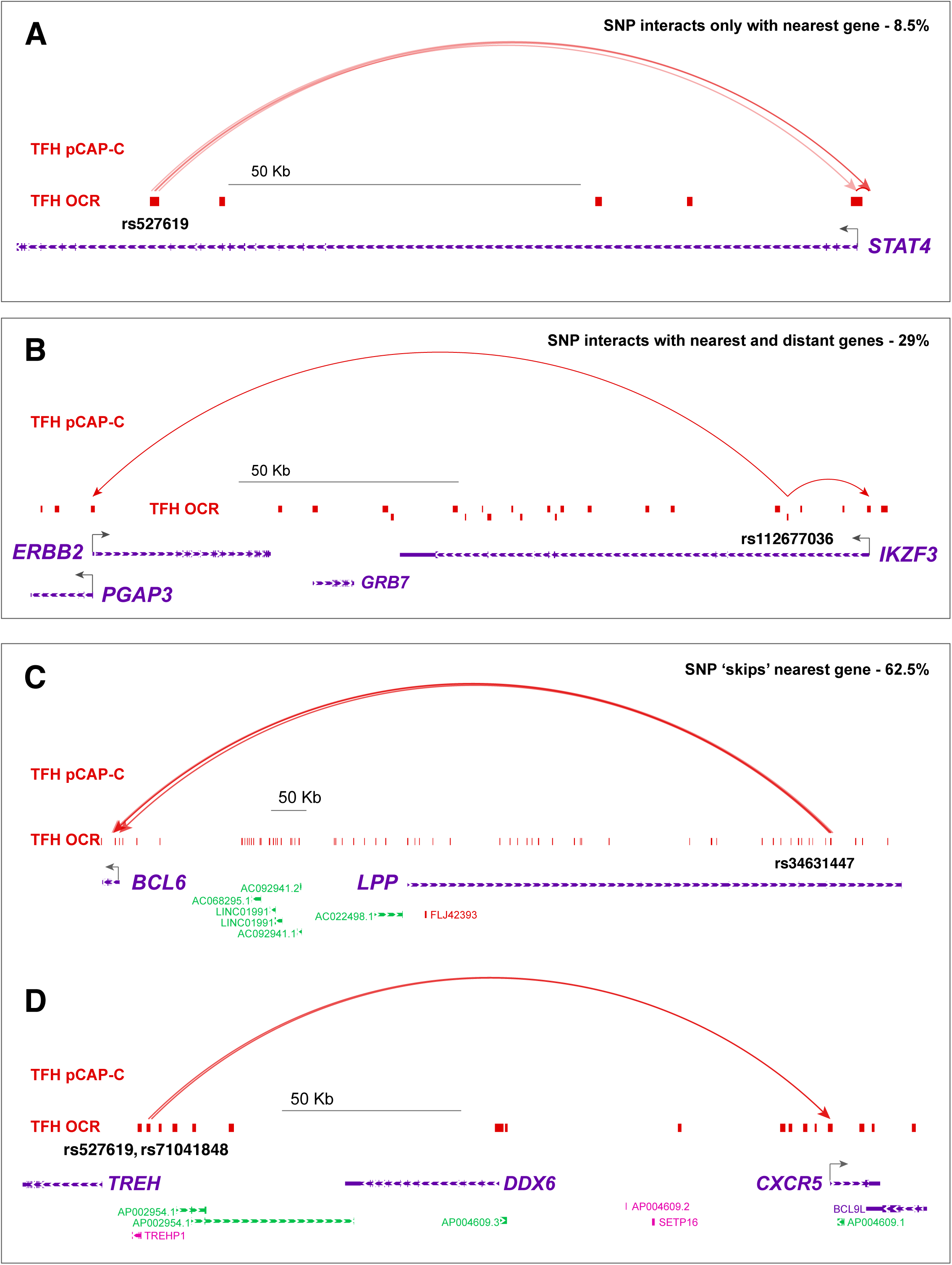
SLE variant-to-gene mapping through integration of GWAS and promoter-open chromatin connectomes in follicular helper T cells. Four categories of accessible SLE SNP-promoter interactions were detected. a. Accessible SLE SNP uniquely interacts with the nearest promoter (8.5%). An example is rs527619, which physically interacts only with the nearest gene STAT4. b. Accessible SLE SNP interacts with the nearest promoter and at least one distant promoter (29%). An example is rs112677036, which physically interacts with IKZF3 and the distant ERBB2 and PGAP3 genes. c. Accessible SLE SNP ‘skips’ the nearest promoter to interact exclusively with on or more distant promoters (62.5%). Examples are (c) rs34631447, which skips LPP to physically interact with BCL6, and (d) rs527619 and rs71041848, which interact with the distant CXCR5 gene instead of the nearest gene, TREH.

Ontology of the set of genes found physically connected to open SLE variants showed enrichment for pathways involved in dendritic cell maturation, T-B cell interactions, T helper differentiation, NFkB signaling, and costimulation through CD28, ICOS, and CD40 (**Figure 7A**). The top three disease networks enriched in SLE SNP-connected genes are systemic autoimmune disorders, rheumatic disease, and type 1 diabetes, all inflammatory disorders involving autoantibody-mediated pathology (**Figure 7B**). At least 200 of these connected genes are differentially expressed between naïve and follicular helper T cells (**Supplemental Tables 2** and **6**), and many have known roles in TFH and/or T cell function (*e.g*., *BCL6, CXCR5, TCF7, PRDM1, IKZF3, IKZF2, IRF8, ETS1, ELF1, EBI3, PTPN22, PDL1, TET3, IL19, IL20*). Similarly, SLE SNP-connected genes are highly regulated (*P*<10^-6^) in a hierarchical manner by IFNg, IL-2, IL-21, IL-1, IL-27, CD40L, and TCR/CD28 (**Figure 7C**). We also compared our list of genes found physically associated with SLE SNPs in TFH with those found statistically associated with SLE variants through eQTL studies in two distinct human subject cohorts. One study by Odhams *et al*. identified 97 gene-SNP eQTL in B-LCL lines^29^, while another by Bentham et al. used B-LCL lines and undifferentiated leukocyte subsets from peripheral blood to identify 41 SLE eQTL^7^. Over one-third (14/41) of the SNP-gene associations implicated by the Bentham study were also identified by our promoter-Capture-C approach in tonsillar TFH cells from three healthy donors (*ANKS1A, C6orf106, RMI2, SOCS1, PXK, UHRF1BP1, LYST, NADSYN1, DHCR7, C15orf39, MPI, CSK, ULK3, FAM219B*; **Supplemental Figure 7**). Similarly, 13% (13/97) of the genes implicated by Odhams *et al.* were found connected to the same SLE SNPs in tonsillar TFH cells (*LYST, NADSYN1, DHCR7, C15orf39, MPI, CSK, ULK3, FAM219B*, *TNIP1, CCDC69, SPRED2, RP11,* **Supplemental Figure 7**), for a total of 16% (19/119) SNP-gene pairs overlapping between both studies. These results indicate that a gene’s spatial proximity to an accessible, disease-associated SNP in 3D is a strong predictor of its role in the context of both normal TFH biology and SLE disease pathogenesis.

**FIGURE 7.**
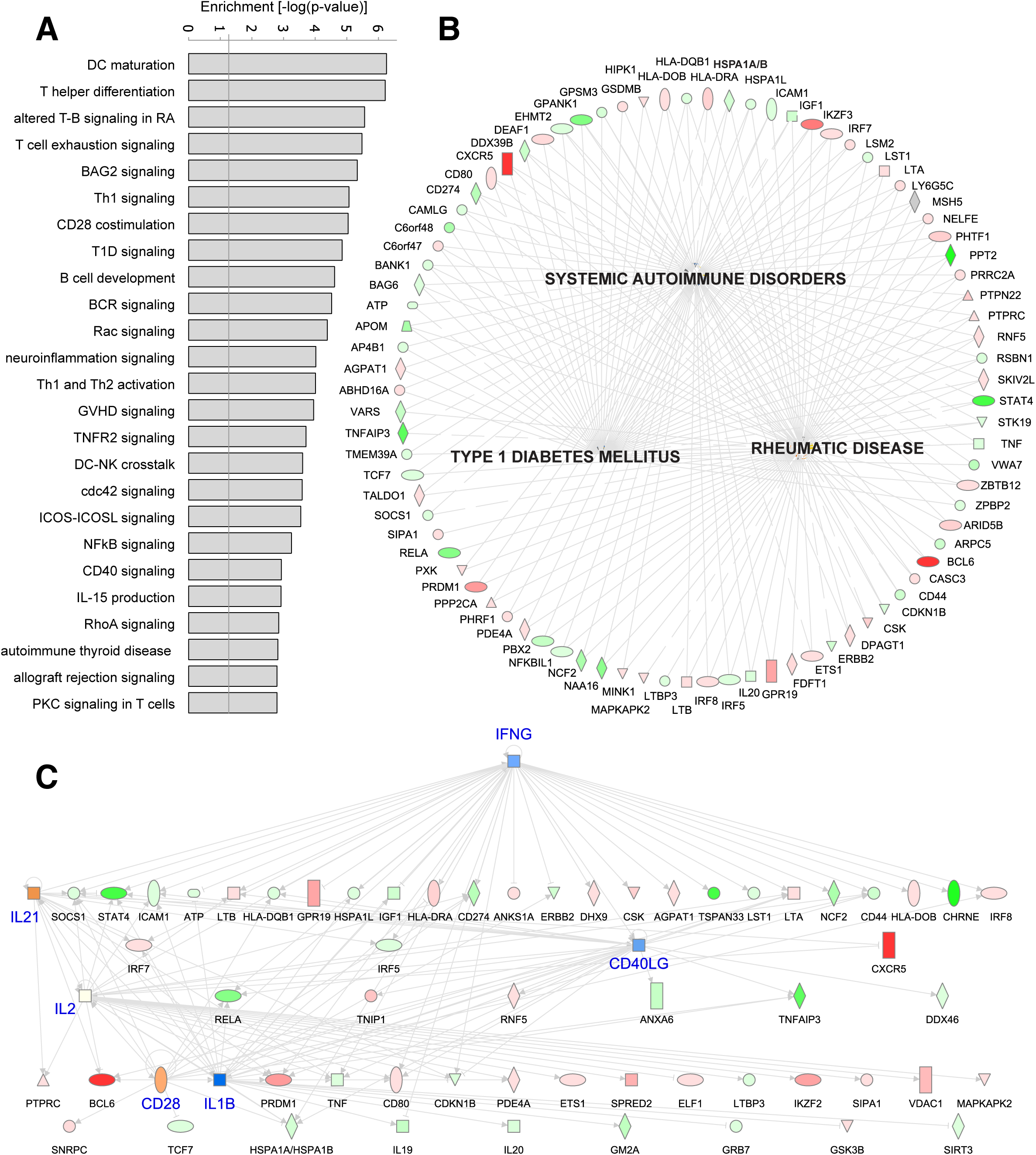
Ontology and pathway analysis of genes implicated through integration of GWAS and promoter-open chromatin connectomes in follicular helper T cells from human tonsil. a. Enrichment of the top 25 canonical pathways (a) and 3 disease networks (b) among genes implicated through promoter-open chromatin connectomes in TFH. c. Regulatory hierarchical network from SLE-connectome-implicated genes. Color gradients in b and c represent log 2 expression changes between TFH and naïve T cells, with green indicating down-regulation and red indicating up-regulation in TFH. Blue nodes in c represent regulatory hubs for genes with no SLE-OCR connectome detected.

### Genomic regions identified by SLE GWAS and 3D epigenomics regulate major TFH genes

To validate that genomic regions implicated by ATAC-seq, promoter-focused Capture-C, and SLE-associated genetic variation function as *bona fide* distal regulatory elements for their connected promoters, we used CRISPR/CAS9 to specifically delete several iOCR harboring SLE variants from the Jurkat T cell genome. We first targeted the intergenic region near the *TREH* gene that harbors the rs527619 and rs71041848 proxies to the rs4639966 SLE sentinel SNP, and was captured interacting with the *CXCR5* promoter (**Supplemental Figure 8**). Neither untargeted parental Jurkat cells nor a control-targeted Jurkat line express CXCR5, but deletion of this region led to induction of CXCR5 in approximately half of the cells (**Figure 8A**). Similarly, parental and control-targeted Jurkat cells do not express *IKZF1*, which encodes the transcription factor Ikaros, but deletion of the OCR containing the rs4385425 proxy SNP to the sentinel SLE SNP rs11185603 (**Supplemental Figure 8**) induced expression of Ikaros in nearly half of the cells (**Figure 8B**). We also targeted the TFH-specific open chromatin region in the sixth intron of the *LPP* gene (**Supplemental Figure 8**) that harbors the rs34631447 and rs79044630 SNPs proxy to sentinel rs6762714 SLE SNP, and was observed interacting with the promoter of *BCL6*. *BCL6* is not expressed by parental or control-targeted Jurkat cells, but is induced by IFN-gamma (**Figure 8C**). However, inducible expression of *BCL6* was completely abrogated in Jurkat cells lacking the ∼150 bp SLE-associated LPP OCR (**Figure 8C**). These results confirm that these distal SLE-associated regions, which are located hundreds to thousands of kilobases away in one dimension, interact with and act as crucial regulatory elements for the genes encoding the master TFH transcription factor *BCL6*, the *IKZF1* transcriptional repressor, and the TFH chemokine receptor *CXCR5*. These results indicate that the 3D promoter connectomes detected in these cells reveal *bona fide* gene regulatory architectures.

**FIGURE 8.**
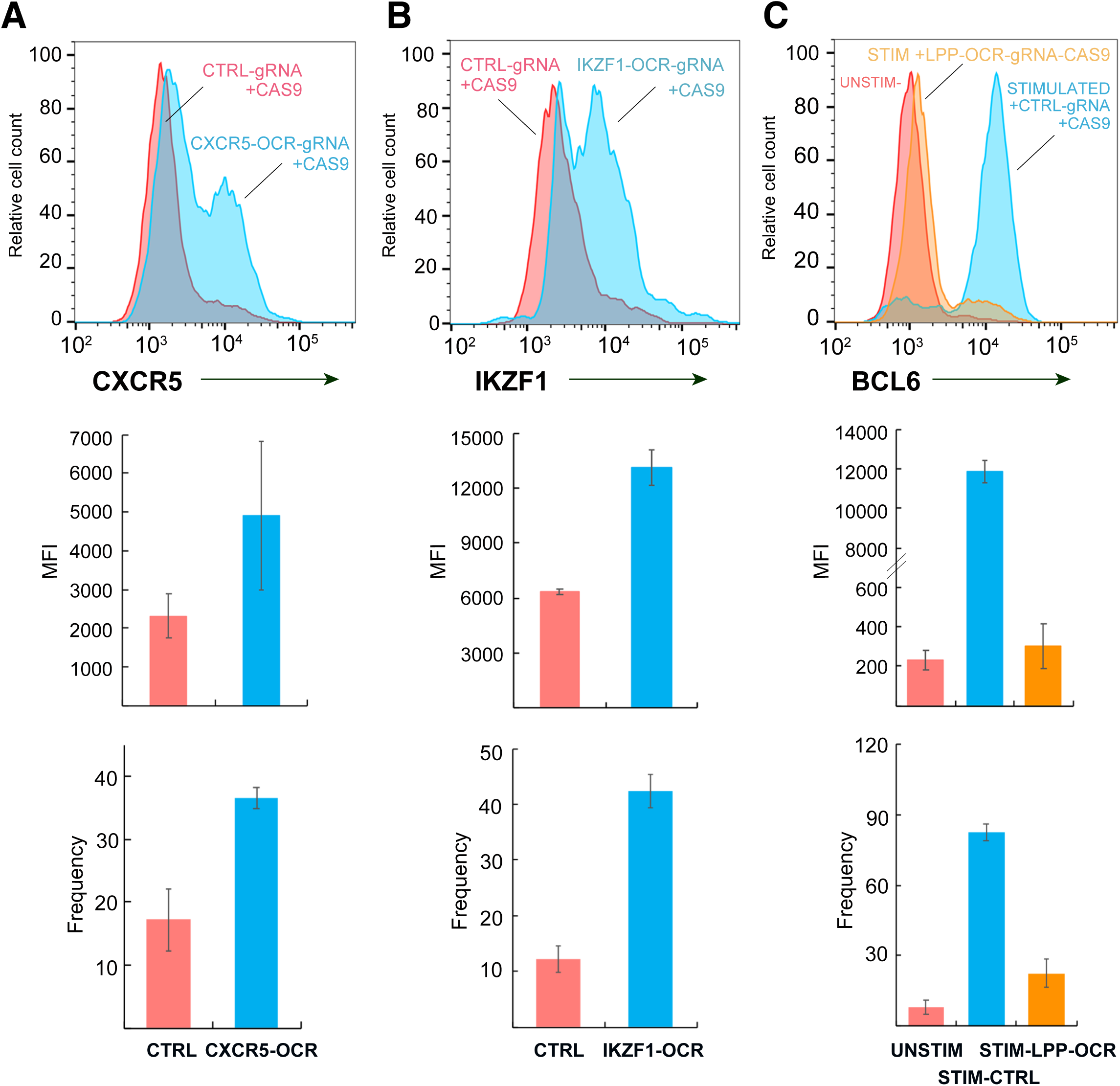
CRISPR/CAS9-based deletion of accessible, promoter-connected genomic regions harboring SLE variants influences TFH gene expression. a, CRISPR-CAS9 targeting of the 136 bp OCR near the TREH gene harboring the rs527619 and rs71041848 SLE proxy SNPs and captured interacting with the CXCR5 promoter leads to increased CXCR5 expression (blue histogram) by Jurkat cells compared to cells transduced with a CTRL-gRNA+CAS9 (pink histogram). b, CRISPR-CAS9 targeting of the 213 bp OCR containing the rs4385425 SLE proxy and captured interacting with the IKZF1 promoter leads to increased IKZF1 (Ikaros) expression (blue histogram) by Jurkat cells compared to cells transduced with a CTRL-gRNA+CAS9 (pink histogram). c, CRISPR-CAS9 targeting of the LPP SLE proxy SNPs rs34631447 and rs79044630 captured interacting with the BCL6 promoter abrogates IFNg-induced BCL-6 expression (orange histogram) compared to cells transduced with a CTRL-gRNA+CAS9 (blue histogram). The red histogram shows BCL-6 expression by unstimulated Jurkat cells. Bar graphs in a-c depict the mean fluorescence intensity (upper panels) and percent positive cells (lower panels) for CXCR5, Ikaros, and BCL-6 in control gRNA-tranduced vs. targeted cells. All data are representative of three independent experiments. See Supplemental Figure 7 for design and validation of CRISPR/CAS9-mediated deletion and mutation.

### SLE-associated open chromatin-promoter connectomes implicate novel genes involved in TFH function

From the set of 243 promoter-connected open SLE variants in TFH cells, we noted a subset of variants that skipped nearby promoters to interact with genes that are upregulated upon TFH differentiation, but have no known specific role in TFH biology. These genes are enriched in canonical pathways such as mannose degradation (*MPI*), epoxysqualene biosynthesis (*FDFT1*), di- and tri-acylglycerol biosynthesis (*LCLAT1, AGPAT1*), cholesterol biosynthesis (*DHCR7, FDFT1*), oxidized GTP/dGTP detoxification (*DDX6*), breast and lung carcinoma signaling (*ERRBB2, HRAS, RASSF5, CDKN1B*), tRNA splicing (*TSEN15, PDE4A*), pentose phosphate pathway (*TALDO1*), acetyl-coA biosynthesis (*PDHB*), dolichyl-diphosphooligosaccharide biosynthesis (*DPAGT1*), and valine degradation (*HIBADH*). Two of these genes, *HIPK1* and *MINK1* (**Figure 9A**), encode a homeobox-interacting kinase and a MAP3/4K homolog that each regulate gene expression in other cell types^30, 31^. Like many genes in this category, both *HIPK1* and *MINK1* are upregulated in TFH, and their promoters interact with OCR that are genetically associated with SLE risk, suggesting they are involved in TFH function. To test this, we transduced TFH differentiated *in vitro* from naïve CD4+ T cells^32^ (**Figure 9B** and **C**) with a lentiviral vector expressing shRNA targeting the *HIPK1* transcript to knock down HIPK1 expression (**Figure 9D**), or with scrambled or *B2M* shRNA as controls (**Supplemental Figure 9**). GFP+ cells were sorted, restimulated with CD3/28 beads, and secretion of IL-21, the major cytokine required for T cell help for B cell antibody production, was measured in the supernatant by ELISA. Remarkably, targeting of HIPK1 expression had no effect on *in vitro* TFH differentiation as measured by induction of BCL6, PD-1 or CXCR5 (**Supplemental Figure 9**), but resulted in a ∼3-fold decrease in IL-21 production (**Figure 9E**). To determine if pharmacologic targeting of HIPK1 can also modulate TFH function, we treated *in vitro* differentiated TFH with the HIPK1/2 inhibitor A64. As with genetic targeting, pharmacologic inhibition of HIPK activity resulted in a dose-dependent reduction in IL-21 production by activated TFH cells (**Figure 9F**) without effecting proliferation, viability, or differentiation (**Supplemental Figure 9**). As a further test of whether SLE-associated promoter-OCR connectomes can reveal novel drug targets for TFH function, we targeted MINK1 pharmacologically with the MAP3/4K antagonist PF06260933. Treatment with this inhibitor resulted in a dose-dependent reduction in IL-21 secretion by TFH cells, with an ED_50_ of 5 nM (**Figure 9G**). Unlike the HIPK1 inhibitor, this MINK1 inhibitor did impact T cell IL-2 production and proliferation, but with an ED_50_ 8- to 10-fold higher than its effect on IL-21 production (**Supplemental Figure 9**). These data show that integrated, 3-dimensional maps of disease-associated genetic variation, open chromatin, and promoter connectomes can lead to *bona fide* novel drug targets that control tissue-specific and SLE-relevant biology.

**FIGURE 9.**
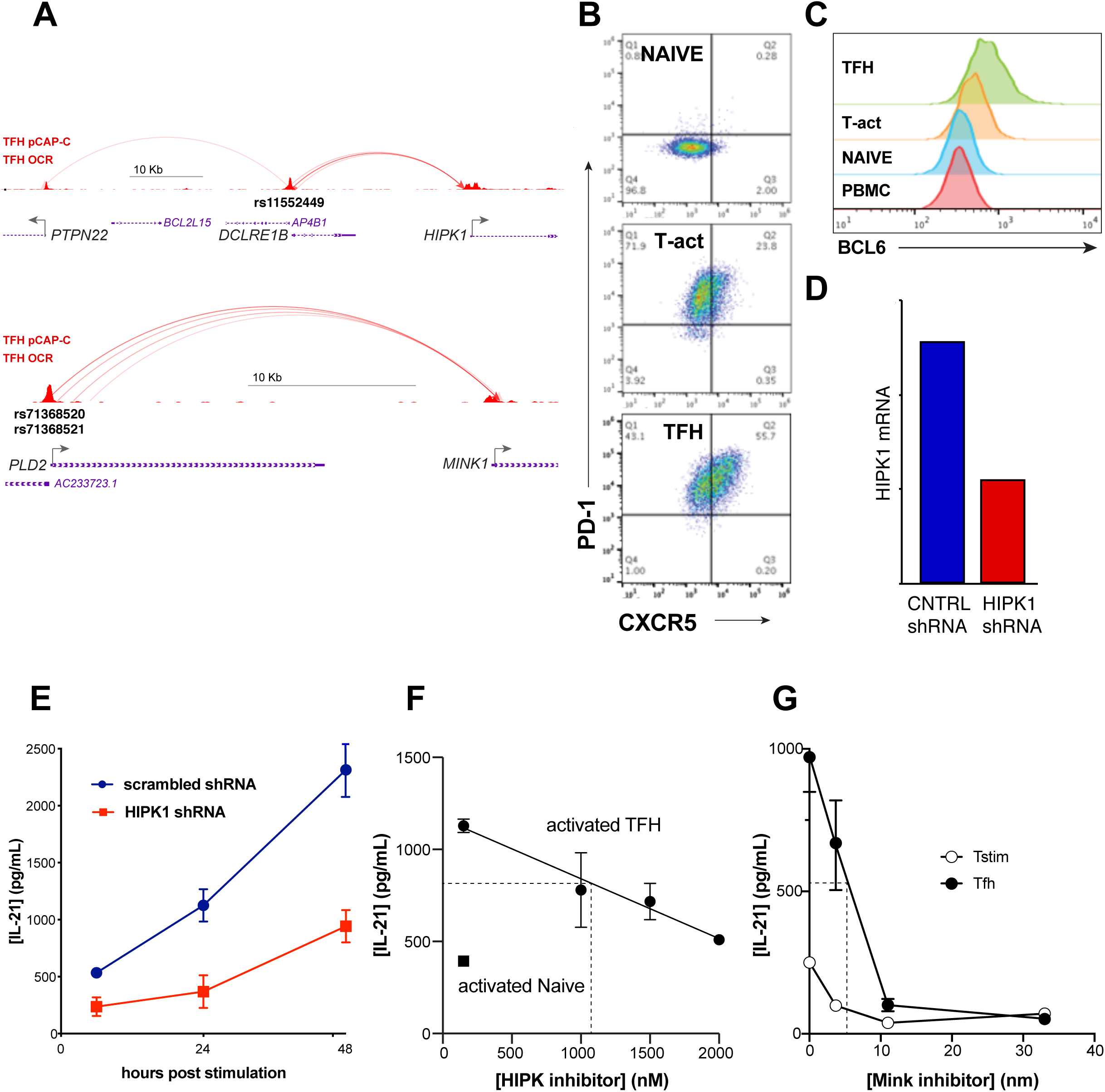
SLE variant-to-gene mapping implicates novel drug targets for modulation of TFH function. a. The interactomes of SLE proxy SNPs rs11552449, rs71368520, and rs71368521 implicate HIPK1 and MINK1. b. Purified naïve CD4+ T cells cultured under TFH-skewing conditions show increased expression of PD-1 and CXCR5, as well as BCL-6 (c). d. HIPK1 mRNA expression by in vitro-differentiated TFH cells transduced with scrambled Lenti-shRNA or Lenti-HIPK-1 shRNA. e. shRNA-mediated knock-down of HIPK1 inhibits IL-21 secretion by TFH cells. f. A HIPK inhibitory drug causes dose-dependent inhibition of IL-21 production by TFH cells. g. A MINK inhibitory drug causes dose-dependent inhibition of IL-21 production by TFH cells. All data are representative of 3-4 independent experiments.

## DISCUSSION

In this study, we used systems-level integration of disease-associated genetic variation and 3-dimensional epigenomic maps of the interactions between open chromatin and promoters in a highly disease-relevant tissue to identify putative disease-associated regulatory regions and the genes they influence. Importantly, this study does not aim to measure the impact of SLE-associated genetic variation on chromatin accessibility or promoter interactions, and therefore does not require large sample sizes from SLE patients and normal individuals. Rather, we use the location of reported, potentially causal SLE variants as ‘signposts’ to identify long-range open chromatin elements that may regulate SLE-relevant gene expression in the context of normal follicular helper T cell biology.

Our one-dimensional open chromatin analyses demonstrated a strong correlation between promoter accessibility and differential gene expression in human tonsillar naive vs. follicular helper T cells. These analyses also showed that SLE-associated variants in accessible promoters in TFH generally coincide with highly expressed genes enriched in autoimmune disease pathways, suggesting that TFH open chromatin landscapes represent a useful filter through which functional, systemic autoimmune disease-associated variants can be identified out of the thousands of sentinel and proxy SNPs to the sentinel signals implicated by GWAS.

However, only 20% of OCR are located in promoter regions, while 80% of the open chromatin regions in human naive and follicular helper T cells map to non-coding/non-promoter regions of the genome, making an interpretation of the potential regulatory role of these regions challenging. To overcome this problem, we generated high-resolution, comprehensive maps of the open chromatin-promoter connectome in naive and TFH cells, allowing physical assignment of non-coding OCR and SNPs to genes, and revealing the potential regulatory architectures of nearly all coding genes and over half of non-coding genes in human immune cell types with crucial roles in humoral immunity and systemic immunopathology. Similar to previous promoter interactome studies^33^, we found that promoter-interacting regions are enriched for open chromatin and the chromatin-based signatures of enhancers. However, we also found that open chromatin regions that interact with a promoter are enriched over 10-fold for enhancer marks compared to OCR that are not connected to a promoter, suggesting that promoter-focused Capture-C preferentially identifies non-coding regions with gene regulatory activity. We also observed enrichment of enhancer marks at open promoters engaged in promoter-promoter interactions compared to promoters not connected to another promoter, suggesting that promoters may synergize in three dimensions in an enhancer-like manner to augment expression of their connected genes^34^.

Our study shows that, similar to previous estimates^35^, less than 10% of promoter interactions exclusively involve the nearest genes. Over 90% of accessible disease variants interact with distant genes, and that over 60% of open variants skip the nearest gene altogether and exclusively interact only with distant genes. Importantly, we were able to validate direct roles for several SLE-associated distal OCR in the regulation of their connected genes (*BCL6, CXCR5, IKZF1*) using CRISPR/CAS9-mediated editing in human T cells, suggesting that SLE-associated genetic variation at distant loci can operate through effects on genes with known roles in TFH and/or SLE biology. A locus control region ∼130 kb upstream of the *BCL6* gene has been defined previously in germinal center B cells^36^, and we also find evidence for usage of this region by human TFH cells at the level of open chromatin, histone enhancer marks, and long-range connectivity to the *BCL6* promoter (**Supplemental Figure 10**). However, we also observe a much more distant ‘stretch’ enhancer within the *LPP* gene in TFH cells, as evidenced by extensive open chromatin, H3K27 acetylation, and H3K4 mono-methylation (**Supplemental Figure 10**). This region shows extensive connectivity with *BCL6* in the 3D architecture of the nucleus, and the 1 Mb distal SLE-associated *BCL6* enhancer validated by genome editing in this study is contained within this *BCL6* stretch enhancer. This enhancer region is occupied in lymphoid cell lines by NFkB/RelA and POU2F2, both transcription factors known to positively regulate immunoglobulin and inflammatory gene expression (ENCODE). Long-range regulatory elements for CXCR5 have not been previously identified, and the −180 kb SLE-associated element in this study is the first validated for *CXCR5*. Deletion of this element led to increased expression of *CXCR5*, suggesting that unlike the distal *BCL6-LPP* enhancer, this element is a silencer in Jurkat cells. Consistent with this finding, this region is occupied by the repressive transcription factors YY1, BHLHE40 and BATF in lymphoid cells (ENCODE), but its function in primary TFH cells remains to be determined. These distant SNP-gene regulatory pairs join examples like the *FTO* and *TCF7L2* loci^1, 3^, in which GWAS data were interpreted to implicate the nearest genes, while 3D epigenomics and functional follow-up showed that the disease variants actually reside in elements that regulate the distant *IRX3* gene (for *FTO*) and the *ACSL5* gene (for *TCF7L2*). Our results indicate that a gene’s spatial proximity in three dimensions to a regulatory SLE SNP is a strong predictor of its function in the context of TFH biology and SLE disease pathogenesis, and suggest that assumptions that a given genomic feature (*e.g.,* SNP or TF binding motif) or epigenomic feature (*e.g*., 5meCpG, 5hmCpG, or histone mark) identified by 1D mapping of the human genome regulates the nearest gene are more likely to be incorrect than correct. These data have important implications for the interpretation of all genetic and epigenomic studies in all tissues.

Remarkably, our integrated open chromatin and promoter connectome mapping in tonsillar TFH cells from three healthy individuals identified one-third of the SNP-gene pairs identified by Bentham^7^, and 13% of the SNP-gene pairs identified by Odhams^29^. These quantitative trait studies require samples from hundreds of individuals, and the data are obtained from blood, B-LCL, or naïve mononuclear leukocytes. However, immune responses do not take place in the blood, and the pathophysiologic aspects of inflammatory disease are mediated by specialized, differentiated immune cell types that are rare or not present in blood. Our approach utilized human follicular helper T cells from tonsil that are ‘caught in the act’ of mediating coordinated *in vivo* T-B immune responses, and is the same cell type involved in B cell help for autoantibody production in SLE. In addition, this variant-to-gene mapping approach identified ∼10-fold more SLE SNP-gene association than current eQTL studies; further follow up work will determine how many of these associations are valid vs. false positives.

In addition to revealing the previously unknown SLE-associated regulatory architectures of known TFH/SLE genes, we show that the combination of GWAS and 3-dimensional epigenomics can identify genes with previously unappreciated roles in disease biology through their connections with accessible disease SNPs. In a previous study, we implicated the novel gene *ING3* by virtue of its interaction with an accessible osteoporosis-associated SNP, and showed that this gene is required for osteoclast differentiation in an *in vitro* model^14^. In this current study, approximately two dozen ‘novel’ genes up- or down-regulated during differentiation of naive CD4+ T cells into TFH were implicated through their connection to SLE SNPs. Among these are *HIPK1*, a nuclear homeobox-interacting protein kinase that cooperates with homeobox, p53, and TGFB/Wnt pathway transcription factors to regulate gene transcription^30, 37–39^. A role for HIPK1 in T-independent B cell responses has be identified in the mouse^40^, but no role for this kinase has been previously established in TFH or SLE. Another gene implicated in our study is *MINK1*, which encodes the misshapen-like kinase MAP4K6. This kinase functions upstream of JNK and SMAD in neurons^41, 42^, and has been shown to inhibit TGFB-induced Th17 differentiation^43^. However, a role in TFH or SLE has likewise not been previously appreciated. We show that genetic and/or pharmacologic targeting of HIPK1 or MINK1 in human TFH cells inhibits their production of IL-21, a cytokine required for T cell-mediated help for B cell antibody production^44^. While further work is required to elucidate the role of these kinases in TFH biology and the pathogenesis of systemic autoimmunity, these examples show the utility of this integrated approach in identifying novel targets for drug repurposing or new compound development in complex heritable diseases.

## METHODS

### Purification of naïve and follicular helper T cells from human tonsil

Fresh tonsils were obtained from immune-competent children (n=10) undergoing tonsillectomy to address airway obstruction or a history of recurrent tonsillitis. The mean age of donors was 5.7 years (range 2-16 years) and 50% were male. Tonsillar mononuclear cells were isolated from tissues by mechanical disruption (tonsils were minced and pressed through a 70 micron cell screen) followed by Ficoll-Paque centrifugation. CD19-positive cells were removed (StemCell) and CD4^+^ T cells were enriched with magnetic beads (Biolegend) prior to sorting naïve T cells (CD4^+^CD45RO^-^) and T follicular helper cells (CD4^+^CD45RO^+^CD25^lo^CXCR5^hi^PD1^hi^) on a MoFlo Astrios EQ (Beckman Coulter).

### Cell fixation

We used standard methods for cell fixation^14^. Briefly, 10^7^ TFH or naïve CD4+ T cells were suspended in 10 mL RPMI + 10% FBS, followed by an additional 270uL of 37% formaldehyde and incubation for 10 min at RT on a platform rocker. The fixation reaction was quenched by the addition of 1.5 mL cold 1M glycine (4°C). Fixed cells were centrifuged at 1000 rpm for 5 min at 4°C and supernatants were removed. The cell pellets were washed in 10 ml cold PBS (4°C) followed by centrifugation as above. Cell pellets were resuspended in 5 ml cold lysis buffer (10 mM Tris pH8, 10 mM NaCl, 0.2% NP-40/Igepal supplemented with a protease inhibitor cocktail). Resuspended cell pellets were incubated for 20 minutes on ice, centrifuged at 1800 rpm, and lysis buffer was removed. Cell pellets were resuspended in 1 mL of fresh lysis buffer, transferred to 1.5 mL Eppendorf tubes, and snap frozen in ethanol/dry ice or liquid nitrogen. Frozen cell pellets were stored at −80°C for 3C library generation.

### 3C library generation

We used standard methods for generation of 3C libraries^14^. For each library, 10^7^ fixed cells were thawed at 37°C, followed by centrifugation at RT for 5 mins at 14,000rpm. The cell pellet was resuspended in 1 mL of dH_2_O supplemented with 5 uL 200X protease inhibitor cocktail, incubated on ice for 10 mins, then centrifuged. Cell pellet was resuspended to a total volume of 650 uL in dH_2_O. 50 uL of cell suspension was set aside for pre-digestion QC, and the remaining sample was divided into 3 tubes. Both pre-digestion controls and samples underwent a pre-digestion incubation in a Thermomixer (BenchMark) with the addition of 0.3%SDS, 1x NEB DpnII restriction buffer, and dH2O for 1hr at 37°C shaking at 1,000rpm. A 1.7% solution of Triton X-100 was added to each tube and shaking was continued for another hour. After pre-digestion incubation, 10 ul of DpnII (NEB, 50 U/µL) was added to each sample tube only, and continued shaking along with pre-digestion control until the end of the day. An additional 10 µL of DpnII was added to each digestion reaction and digested overnight. The next day, a further 10 µL DpnII was added and continue shaking for another 2-3 hours. 100 uL of each digestion reaction was then removed, pooled into one 1.5 mL tube, and set aside for digestion efficiency QC. The remaining samples were heat inactivated incubated at 1000 rpm in a MultiTherm for 20 min at 65°C to inactivate the DpnII, and cooled on ice for 20 additional minutes. Digested samples were ligated with 8 uL of T4 DNA ligase (HC ThermoFisher, 30 U/µL) and 1X ligase buffer at 1,000 rpm overnight at 16°C in a MultiTherm. The next day, an additional 2 µL of T4 DNA ligase was spiked in to each sample and incubated for another few hours. The ligated samples were then de-crosslinked overnight at 65°C with Proteinase K (20 mg/mL, Denville Scientific) along with pre-digestion and digestion control. The following morning, both controls and ligated samples were incubated for 30 min at 37°C with RNase A (Millipore), followed by phenol/chloroform extraction, ethanol precipitation at −20°C, the 3C libraries were centrifuged at 3000 rpm for 45 min at 4°C to pellet the samples. The controls were centrifuged at 14,000 rpm. The pellets were resuspended in 70% ethanol and centrifuged as described above. The pellets of 3C libraries and controls were resuspended in 300uL and 20μL dH_2_O, respectively, and stored at −20°C. Sample concentrations were measured by Qubit. Digestion and ligation efficiencies were assessed by gel electrophoresis on a 0.9% agarose gel and also by quantitative PCR (SYBR green, Thermo Fisher).

### Promoter-Capture-C design

Our promoter-Capture-C approach was designed to leverage the four-cutter restriction enzyme *DpnII* in order to give high resolution restriction fragments of a median of ∼250bp^14^. This approach also allows for scalable resolution through *in silico* fragment concatenation (**Supplemental Table 4**). Custom capture baits were designed using Agilent SureSelect RNA probes targeting both ends of the *DpnII* restriction fragments containing promoters for coding mRNA, non-coding RNA, antisense RNA, snRNA, miRNA, snoRNA, and lincRNA transcripts (UCSC lincRNA transcripts and sno/miRNA under GRCh37/hg19 assembly) totaling 36,691 RNA baited fragments through the genome^14^. In this study, the capture library was re-annotated under gencodeV19 at both 1-fragment and 4-fragment resolution, and is successful in capturing 89% of all coding genes and 57% of noncoding RNA gene types. The missing coding genes could not be targeted due to duplication or highly repetitive DNA sequences in their promoter regions.

### Promoter-Capture-C assay

Isolated DNA from 3C libraries was quantified using a Qubit fluorometer (Life technologies), and 10 μg of each library was sheared in dH_2_O using a QSonica Q800R to an average fragment size of 350bp. QSonica settings used were 60% amplitude, 30s on, 30s off, 2 min intervals, for a total of 5 intervals at 4 °C. After shearing, DNA was purified using AMPureXP beads (Agencourt). DNA size was assessed on a Bioanalyzer 2100 using a DNA 1000 Chip (Agilent) and DNA concentration was checked via Qubit. SureSelect XT library prep kits (Agilent) were used to repair DNA ends and for adaptor ligation following the manufacturer protocol. Excess adaptors were removed using AMPureXP beads. Size and concentration were checked by Bioanalyzer using a DNA 1000 Chip and by Qubit fluorometer before hybridization. One microgram of adaptor-ligated library was used as input for the SureSelect XT capture kit using manufacturer protocol and our custom-designed 41K promoter Capture-C library. The quantity and quality of the captured library was assessed by Bioanalyzer using a high sensitivity DNA Chip and by Qubit fluorometer. SureSelect XT libraries were then paired-end sequenced on 8 lanes of Illumina Hiseq 4000 platform (100 bp read length).

### ATAC-seq library generation

A total of 50,000 to 100,000 sorted tonsillar naive or follicular helper T cells were centrifuged at 550g for 5 min at 4°C. The cell pellet was washed with cold PBS and resuspended in 50 μL cold lysis buffer (10 mM Tris-HCl, pH 7.4, 10 mM NaCl, 3 mM MgCl2, 0.1% NP-40/IGEPAL CA-630) and immediately centrifuged at 550g for 10 min at 4°C. Nuclei were resuspended in the Nextera transposition reaction mix (25 ul 2x TD Buffer, 2.5 uL Nextera Tn5 transposase (Illumina Cat #FC-121-1030), and 22.5 ul nuclease free H_2_O) on ice, then incubated for 45 min at 37°C. The tagmented DNA was then purified using the Qiagen MinElute kit eluted with 10.5 μL Elution Buffer (EB). Ten microliters of purified tagmented DNA was PCR amplified using Nextera primers for 12 cycles to generate each library. PCR reaction was subsequently cleaned up using 1.5x AMPureXP beads (Agencourt), and concentrations were measured by Qubit. Libraries were paired-end sequenced on the Illumina HiSeq 4000 platform (100 bp read length).

### ATAC-seq analysis

TFH and naïve ATAC-seq peaks were called using the ENCODE ATAC-seq pipeline (https://www.encodeproject.org/atac-seq/). Briefly, pair-end reads from three biological replicates for each cell type were aligned to hg19 genome using bowtie2, and duplicate reads were removed from the alignment. Narrow peaks were called independently for each replicate using macs2 (-p 0.01 --nomodel --shift -75 --extsize 150 -B --SPMR --keep-dup all --call-summits) and ENCODE blacklist regions (ENCSR636HFF) were removed from peaks in individual replicates. Peaks from all replicates were merged by bedtools (v2.25.0) within each cell type and the merged peaks present in less than two biological replicates were removed from further analysis. Finally, ATAC-seq peaks from both cell types were merged to obtain reference open chromatin regions. To determine whether an OCR is present in TFH and/or naïve cells, we first intersected peaks identified from individual replicates in each cell type with reference OCRs. If any peaks from at least one replicate overlapped with a given reference OCR, we consider that region is open in the originating cell type. Quantitative comparisons of TFH and naïve open chromatin landscapes were performed by evaluating read count differences against the reference OCR set. De-duplicated read counts for OCR were calculated for each library and normalized against background (10K bins of genome) using the R package csaw (v 1.8.1). OCR peaks with less than 1.5 CPM (4.5 ∼ 7.5 reads) support in the top 50% of samples were removed from further differential analysis. Differential analysis was performed independently using edgeR (v 3.16.5) and limmaVoom (v 3.30.13). Differential OCR between cell types were called if FDR<0.05 and absolute log2 fold change >1 in both methods.

### Promoter-focused Capture-C analysis

Paired-end reads from three biological replicates for naïve and follicular helper T cells were pre-processed using the HICUP pipeline (v0.5.9) {Wingett:2015}, with bowtie2 as aligner and hg19 as the reference genome. We were able to detect non-hybrid reads from all targeted promoters, validating the success of the promoter capture procedure. Significant promoter interactions at 1-DpnII fragment resolution were called using CHiCAGO (v1.1.8) {Cairns:2016} with default parameters except for binsize set to 2500. Significant interactions at 4-DpnII fragment resolution were also called using CHiCAGO with artificial .baitmap and .rmap files in which DpnII fragments were concatenated *in silico* into 4 consecutive fragments using default parameters except for removeAdjacent set to False. The significant interactions (CHiCAGO score > 5) from both 1-fragment and 4-fragment resolutions were exported in .ibed format and merged into a single file using custom a PERL script to remove redundant interactions and to keep the max CHiCAGO score for each interaction. Open chromatin interaction landscapes were established by projecting significant DpnII fragment interactions at merged 1- and 4-fragment resolutions to reference OCR (**Figure 3A**). First, DpnII fragments involved in significant interactions (both “bait” and “other end”) were intersected with reference OCR using bedtools (v2.25.0). Interactions between bait and other end OCR pairs were called independently for each cell type if their overlapped fragments interacted at either resolution and if both OCR were called as “open” in the corresponding cell type. OCR involved in promoter interactions (iOCR) were classified as promoter OCR (prOCR) or regulatory OCR (nonprOCR) by comparing their genomic locations to pre-defined promoter regions (− 1500bp ∼ 500bp of TSS) of transcripts in GENCODE V19 and UCSC noncoding RNA described above. If both ends of an OCR interaction failed to overlap a gene promoter, that interaction was removed. OCR pair interactions were combined from both cell types to obtain the reference open chromatin promoter-captured interaction landscapes.

### Microarray analysis of gene expression

RNA from two biological naïve tonsillar CD4+ T cell replicates and four biological tonsillar TFH replicates were hybridized to Affymetrix Human Clarion S arrays at the CHOP Nucleic Acid and Protein Core. Data were pre-processed (RMA normalization), and analyzed for differential expression (DE) using Transcriptome Analysis Console v 4.0 with a false discovery rate (FDR) threshold of 0.05 and a fold-change (FC) threshold of 2. Lists of differentially expressed genes were generated and ranked by log2 fold change. The log2 fold change of the genes with significantly differential accessibility at promoter regions were compared to the pre-ranked gene expression data for GSEA enrichment analysis.

### Gene set enrichment and Ingenuity pathway analysis

Histone mark and CTCF ChIP-seq datasets for naïve and follicular helper T cells were obtained from public resources^19–21^ and compared to promoter-interacting fragments or promoter-interacting OCR. Enrichment of promoter-interacted fragments (PIR) for histone marks and CTCF regions was determined independently in each cell type using the function peakEnrichment4Features() in the CHiCAGO package, and feature enrichment at promoter-interacting OCR were compared to enrichment at non-promoter-interacting OCR using the feature enrichment R package LOLA (v1.4.0)^45^. Fisher’s exact tests were performed and odd ratios were plotted for significant enrichment (pvalue<10^-6^) using ggplot2. The chromatin states of promoter-interacting OCR were also determined using ChromHMM (v1.17) on binarized bed file of histone marks ChIP-seq peaks with 15 states for naïve T cells and 6 states for TFH cells. The annotation of chromatin states was manually added with the reference to epigenome roadmap project^20^. Ingenuity pathway analysis (IPA, QIAGEN) was used for all the pathway analysis. The top significantly enriched canonical pathways were plotted using ggplot2 and networks with relevant genes were directly exported from IPA.

### CRISPR/CAS9 genome editing

CRISPR guide RNAs (sgRNA) targeting rs34631447, rs79044630, rs527619, rs71041848, and rs4385425 were designed using http://crispr.tefor.net and cloned into lentiCRISPRv2-puro or lentiCRISPRv2-mCherry (Feng Zhang, Addgene plasmid #52961; http://n2t.net/addgene:52961; RRID:Addgene_52961) by golden gate ligation using the *BsmB1* restriction enzyme (NEB). 293T cells were transfected in DMEM using Lipofectamine 2000 (Invitrogen) with 6 ug PsPAX2 and 3.5 ug PmD2.G packaging plasmids and 10 ug empty lentiCRISPRv2 or 10 ug sgRNA-encoding lentiCRISPRv2. Viral supernatants were collected after 48 hrs for transduction into Jurkat leukemic T cells maintained in RPMI 1640 with 10% fetal bovine serum, L-glutamine, 2-mercaptoethanol, and penicillin/streptomycin. Cells were seeded in a 24 well plate at 0.5 x10^6^ in 0.5 mL of media per well, and 1 mL of viral supernatant with 8 ug/mL of polybrene was added to each well. Spin-fection was performed for 90 min. at 2500 rpm and 25°C, and transduced cells were equilibrated at 37C for 6 hrs. For rs34631447, rs79044630, and rs4385425, 1.2 ml of media was removed and replaced with 1 ml of fresh media containing 1 ug of puromycin for 7 days of selection before use in experiments. Cells transduced with sgRNAs targeting rs527619 and rs71041848 were sorted based on mCherry on a FACS Jazz (BD Biosciences). Mutations were analyzed by PCR coupled with Sanger sequencing at the CHOP Nucleic Acids and Protein Core. The following primers were used for PCR: BLC6-F: CTCTGTGGTTGTGGGCAAGGC-R:CAGGTGGCGAATCAGGACAGG, CXCR5-F: GTCCCTGGTGATGGAAACTCAGGC-R: GCAGTGGCCTCCCTTACACAGG, IKZF1-F: CCTTCTCCATGCCCAGGTGACTC-R: GGCCTCAGCTAGGCAAACCAGAG. Measurement of BCL-6 expression in targeted Jurkat lines was assessed by flow cytometry using anti-human APC-BCL-6 (Biolegend) after treatment with human recombinant IFNg (5 ng/mL, R&D Systems) overnight and stimulation with PMA (30 ng/mL) and ionomycin (1 uM, Sigma-Aldrich) for 4-6 hrs. Expression of Ikaros and CXCR5 by targeted Jurkat lines was also assessed by flow cytometry using anti-human APC-CXCR5 (Biolegend) and anti-human PE-Ikaros (BD Biosciences). Fixation, permeabilization and intracellular staining for Ikaros and BCL-6 was performed using the Transcription Factor Buffer Set (BD Pharmingen). Cells were analyzed on a CytoFLEX flow cytometer (Beckman Coulter).

### Lentiviral shRNA-based gene targeting

A lentiviral shRNA-based approach was employed to silence the expression of HIPK1 as well as B2M as a positive control. The lenti-shRNA vectors pGFP-C-shRNA-Lenti-Hipk1, pGFP-C-shRNA-Lenti-B2M and pGFP-C-scrambled were purchased from Origene. The packaging vectors PmD2G and PsPAX.2 were obtained from Addgene. Exponentially growing 293T cells were split and seeded at 8 x 10^6^ cells in 100 mm dishes in RPMI 1640 medium at 37C. The following day, cells were transfected in antibiotic- and serum-free medium with lenti shRNA plus packaging vector DNA prepared in a complex with Lipofectamine 2000. After 6 hrs of transfection, medium was replaced with complete serum containing RPMI medium and cells were cultured at 37C for 2 days. Human primary CD4+ T cells from healthy donors were obtained from the University of Pennsylvania Human Immunology Core and stimulated overnight with human anti-CD3- and anti-CD28-coated microbeads. Cells were harvested, de-beaded, washed with warm RPMI medium, and aliquots of 10^6^ activated CD4+T cells were infected with 1 ml of viral supernatant collected from lenti-shRNA transfected 293T cell cultures. Polybrene was added to the viral supernatant at 8 ug/ml, cells were spin-fected at 2500 rpm for 1.5 hrs, cultured at 37C for 6 hrs, and restimulated with anti-CD3 and anti-CD28 beads, Activin A (100 ng/ml), IL-12 (5 ng/ml), and anti-IL-2 (2 ug/ml) to induce *in vitro* TFH differentiation {Locci;2016}. After 4 days of differentiation, transduced cells were FACS-sorted based on GFP expression, and expression of B2M, BCL-6, CXCR5 and PD-1 was measured by flow cytometry. In addition, sorted GFP+ *in vitro* TFH cells were restimulated with plate-bound human anti-CD3 and anti-CD28 (1 ug/ml each) in flat bottom 96 well plates, and supernatants were collected at the indicated timepoints for assessment of IL-21 secretion by ELISA. RNA was extracted from sorted GFP+ TFH cells using an RNeasy micro kit (Qiagen), treated with DNase, and 500 ng of total RNA was reverse-transcribed using iScript cDNA synthesis kit (Bio-Rad). qRT-PCR quantification of HIPK-1, B2M and 18s rRNA transcripts was performed using Amplitaq Gold SYBR Master mix (ABI) on Applied Biosytems step one plus real-time thermocycler. Specific mRNA levels were determined as ratio of total 18s rRNA. The following primer sequences were used for qRT PCR: HIPK-1-F: CAGTCAGGAGTTCTCACGCA, HIPK-1-R: TGGCTACTTGAGGGTGGAGA, B2M-F: GCCGTGTGAACCATGTGACT, B2M-R: CATCCAATCCAAATGCGGCA, hu 18S-F: CCTTTAACGAGGATCCATTGGA, hu 18S-R: CGCTATTGGAGCTGGAATTACC.

### Pharmacologic inhibitors

The HIPK kinase family inhibitor A64 trifluoroacetate was purchased from Sigma, and the MAP4K2 inhibitor PF06260933, which also inhibits MINK1 and TNIK, was purchased from TOCRIS. Human primary CD4+ T cells were cultured under TFH condition for 5 days in the presence of the indicated concentrations of each inhibitor (150 nM to 2500 nM for A64, 3.7 nM to 100 nM for PF06260933). In addition, anti-CD3- and anti-CD28-stimulated human CD4+ T cells (non-TFH) were cultured in the presence of inhibitors. After 5 days of primary culture, cells were harvested and 10^6^ cells were restimulated with plate-bound human anti-CD3+ anti-CD28 (1 ug/ml each) in the presence of inhibitors. Culture supernatants were collected at the indicated timepoints for measurement of IL-2 and IL-21 by ELISA (eBioscience).

### Data availability

Our data are available from ArrayExpress (https://www.ebi.ac.uk/arrayexpress/) with accession numbers E-MTAB-6621 (promoter-Capture-C), E-MTAB-6617 (ATAC-seq), and E-MTAB-6637(expression microarray) respectively.

---

**SUPPLEMENTAL TABLE 1**. Reference ATAC-seq peaks in naive and TFH cells.

**SUPPLEMENTAL TABLE 2**. Microarray-based gene expresssion analysis.

**SUPPLEMENTAL TABLE 3**. OCR harboring SLE proxy SNPs.

**SUPPLEMENTAL TABLE 4**. Summary of 1- and 4-fragment-based promoter interaction calls.

**SUPPLEMENTAL TABLE 5**. Summary of OCR-based promoter interaction calls.

**SUPPLEMENTAL TABLE 6**. SLE variant-to-gene mapping using promoter-OCR connectomes in follicular helper T cells.

## Acknowledgements

This work was supported by The Center for Spatial and Functional Genomics at The Children’s Hospital of Philadelphia (ADW and SFG), the Daniel B. Burke Endowed Chair for Diabetes Research (SFG), the Jeffrey Modell Foundation (NR), and NIH grants AI123539 (ADW), DK122586 (ADW, SFG), HG010067 (SFG), AI146026 (NR), and AI115001 (NR).

## Author Contributions

C.S. assisted with preparation of the manuscript and conducted the epigenomic and transcriptomic analyses with the help of E.M.; A.C. designed the custom Capture-C promoter probe set; C.L.C. provided human tonsillar T cells, N.R. contributed to the design of the immunologic studies and provided microarray data; M.E.J, M.E.L, S.L., K.M.H, and J.P. generated and sequenced the epigenomic libraries; A.T., R.M.T. and P.S. performed the CRISPR/CAS, shRNA and pharmacologic targeting experiments; S.F.G. directed the genomic and epigenomic studies; A.D.W. directed the epigenomic and immunologic studies and wrote the manuscript. C.S., M.E.J., A.T., R.M.T contributed equally, and S.F.G and A.D.W. are co-senior authors.

---

**SUPPLEMENTARY FIGURE 1.**
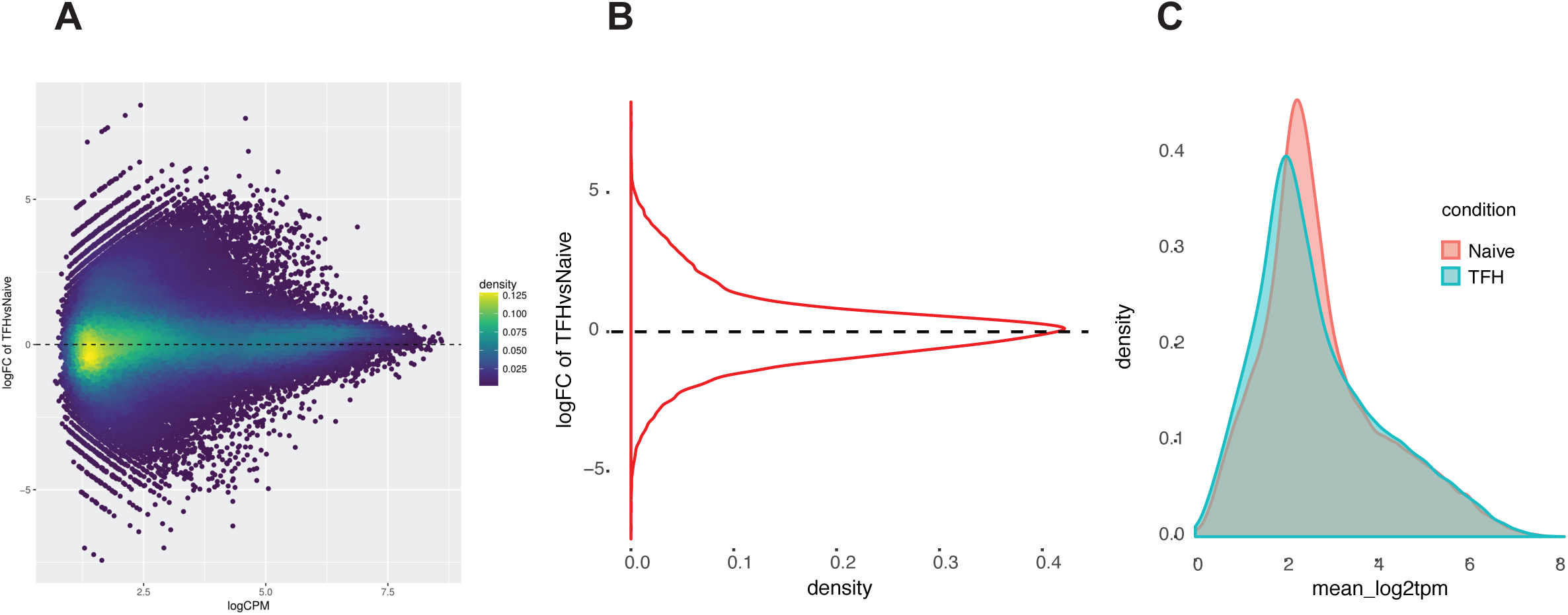
TFH and naive T cells show comparable genomic accessibility. Overall log2 fold changes in reference OCR accessibility (CPM) in TFH compared to naive T cells represented by density plot (a) or distribution plot (b). c. The accessibility signal was normalized by the counts per million method and mean p values across three replicates were used for comparison between TFH and naive T cells.

**SUPPLEMENTARY FIGURE 2.**
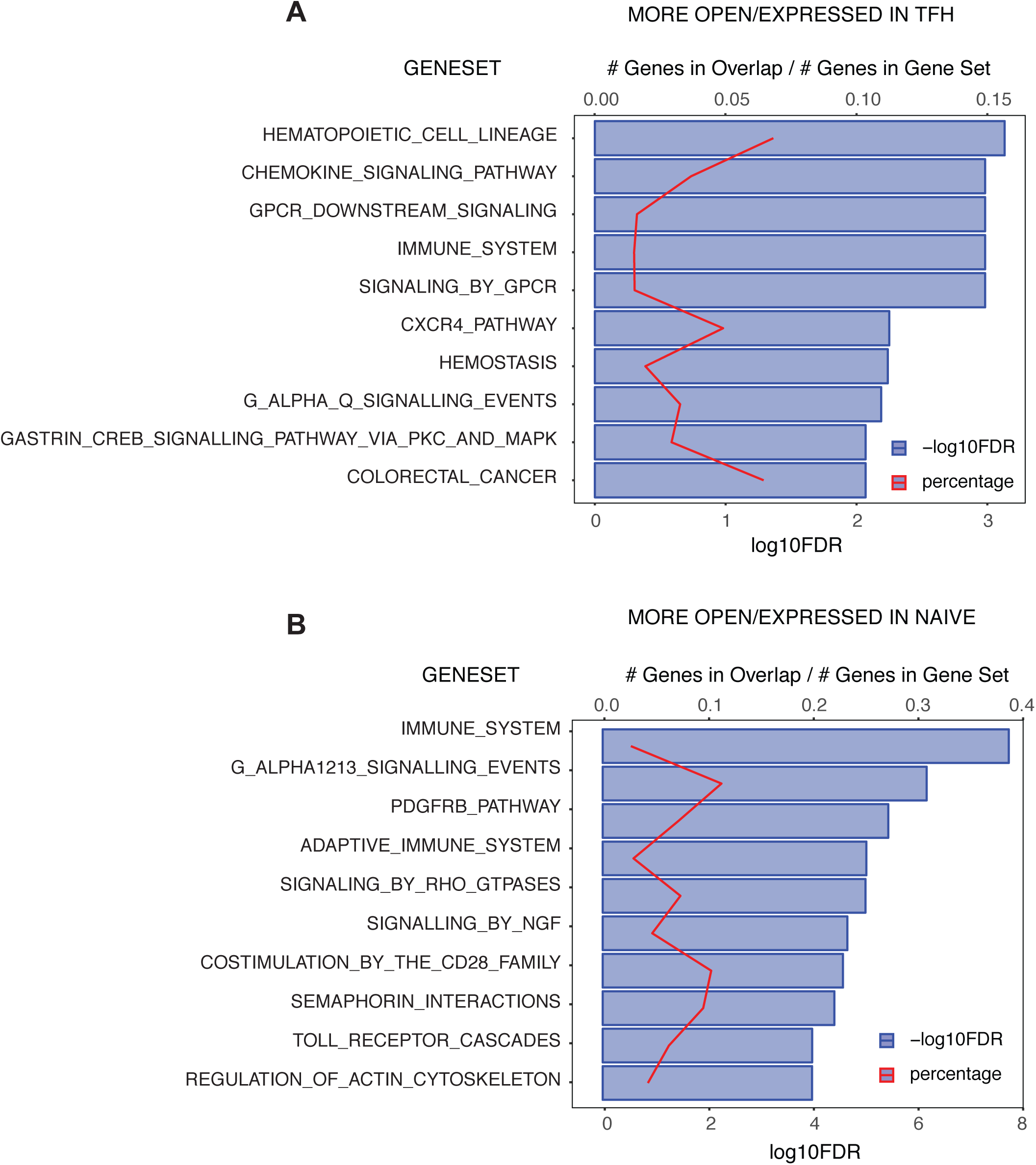
Canonical pathway enrichment for genes with accessible SLE SNPs in their promoters. The log2FDR (blue) and gene ratios (red) for the top 10 enriched Ingenuity canonical pathways is shown for TFH (a) and naive (b) cells.

**SUPPLEMENTARY FIGURE 3.**
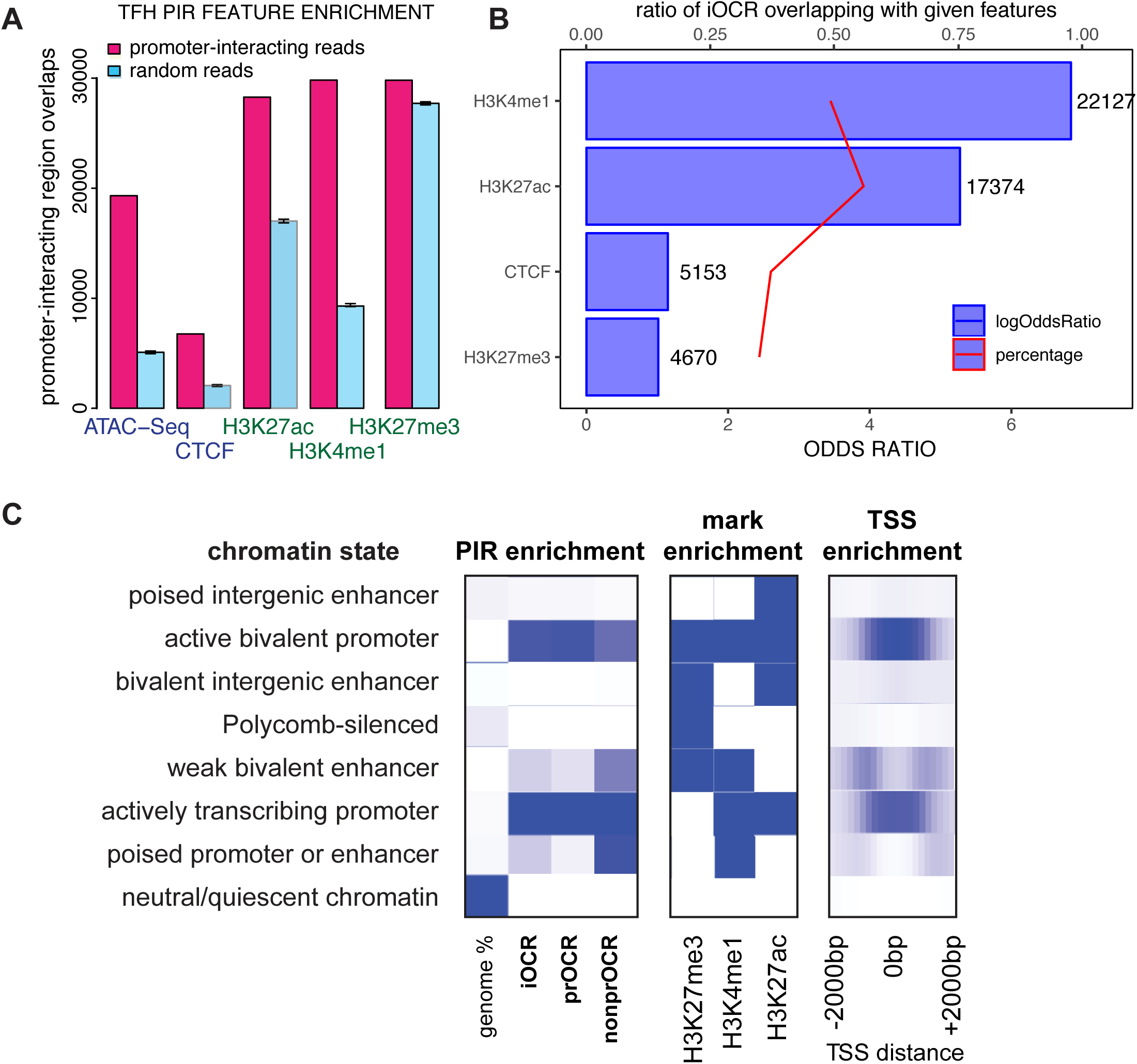
Enrichment of chromatin signatures at promoter interacting regions in TFH cells. a. PIR enrichment for genomic features compared with distance-matched random regions in TFH cells. Error bars show SD across 100 draws of non-significant interactions. b. Feature enrichment of promoter-interacting OCR (iOCR) compared to a random sample of non-promoter-interacting OCR in TFH. c. Enrichment of iOCR within chromHMM-defined chromatin states and TSS neighborhood in TFH. Roadmap Epigenomics 15-state models (middle panel) were defined on the basis of 5 histone modifications (H3K4me1, H3K4me3, H3K27me3, H3K27ac and H3K36me3). Blue color intensity represents the probability of observing the mark in the state. The heatmap to the left of the emission parameters displays the overlap fold enrichment for different categories of iOCR, while the heatmap to the right shows the fold enrichment for each state within 2 kb around a set of TSS. Blue color intensity represents fold enrichment.

**SUPPLEMENTARY FIGURE 4.**
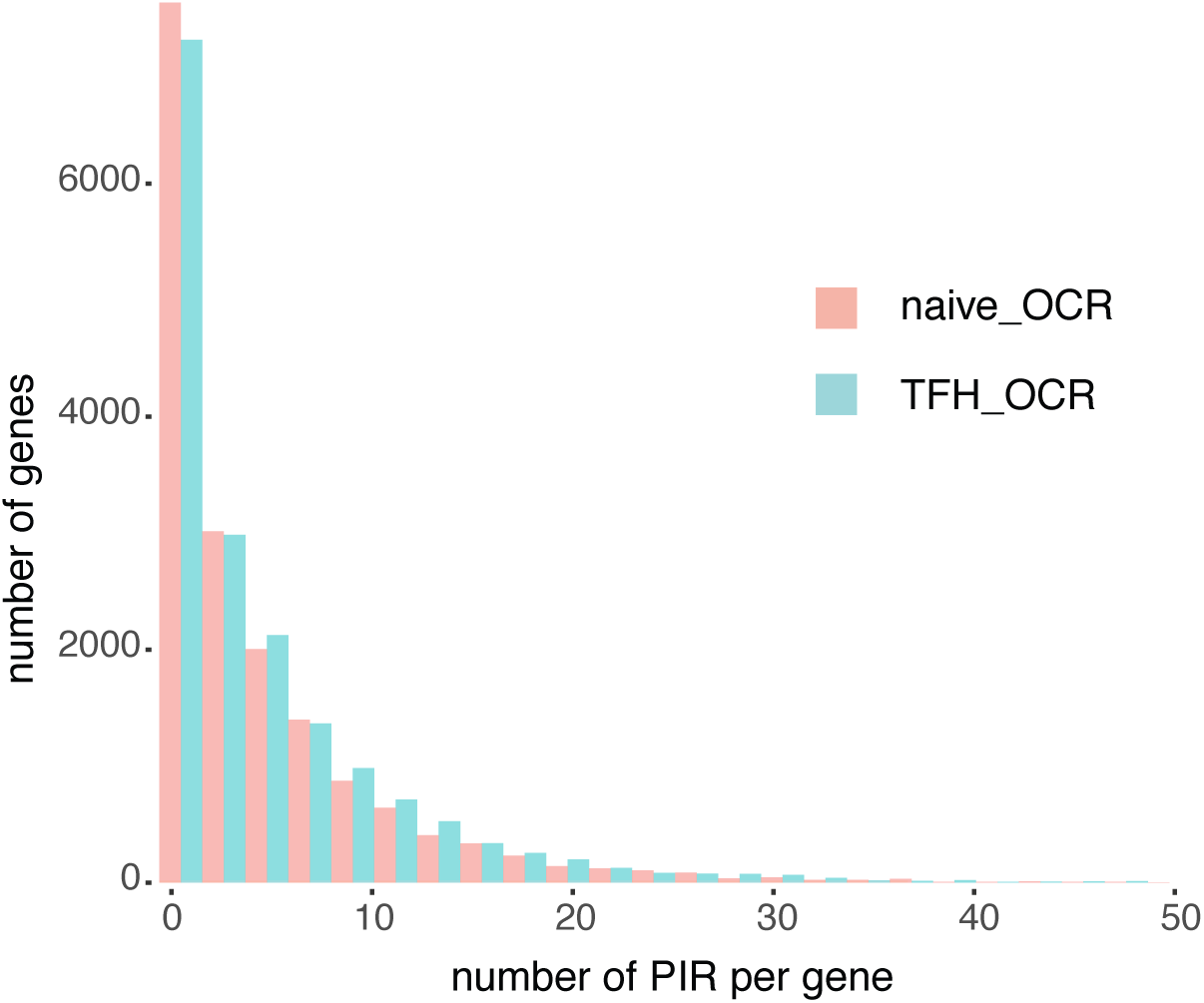
Distribution of promoter-interacting OCR per gene in naïve T and TFH cells. The number of promoter-interacting OCR per gene is plotted for both naïve T (red) and TFH (blue) cells.

**SUPPLEMENTARY FIGURE 5.**
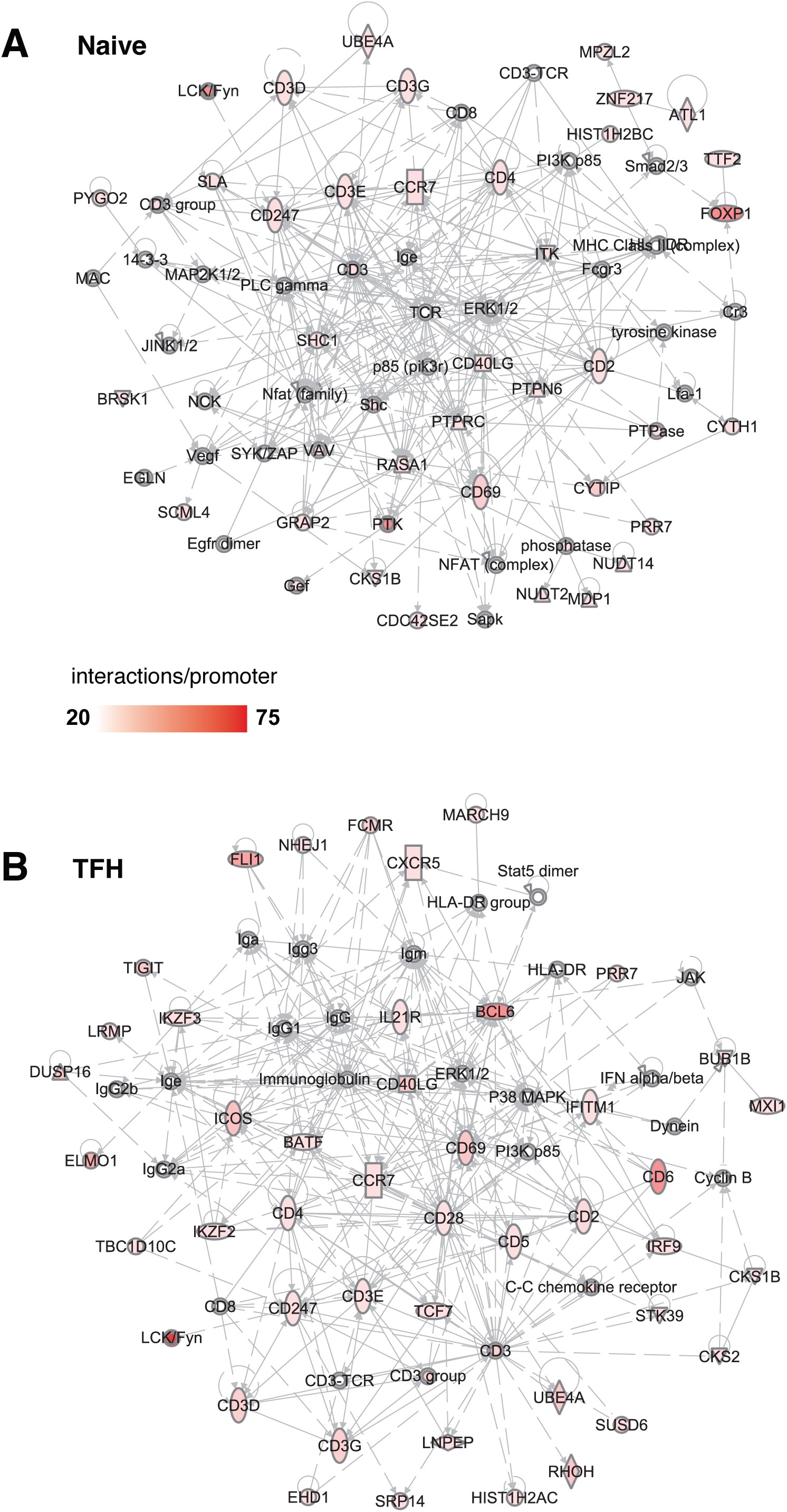
Immune networks enriched among SLE SNP connectome implicated gene sets. The top 3 merged immune networks in naïve (a) and TFH (b) are depicted. Red color intensity represents the number of interactions detected per promoter for each gene in the network.

**SUPPLEMENTARY FIGURE 6.**
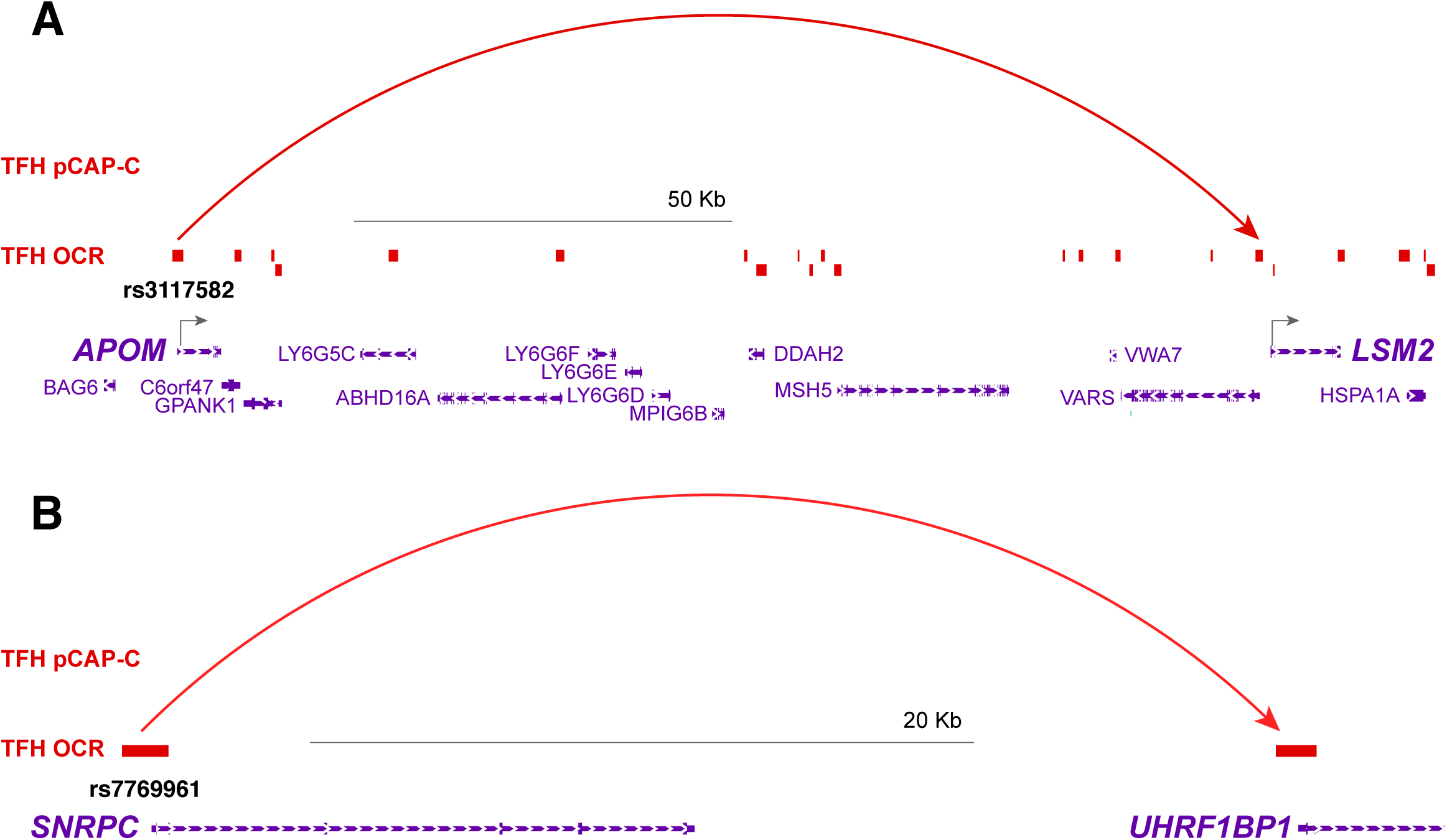
Interaction of open SLE variants with genes encoding nuclear proteins targeted by autoantibodies in SLE patients. a. The accessible SNP rs3117582 at the promoter of APOM physically interacts with the LSM2 promoter. b. The accessible SNP rs7769961 at the SNRPC promoter physically interacts with the UHRF1BP1 promoter.

**SUPPLEMENTARY FIGURE 7.**
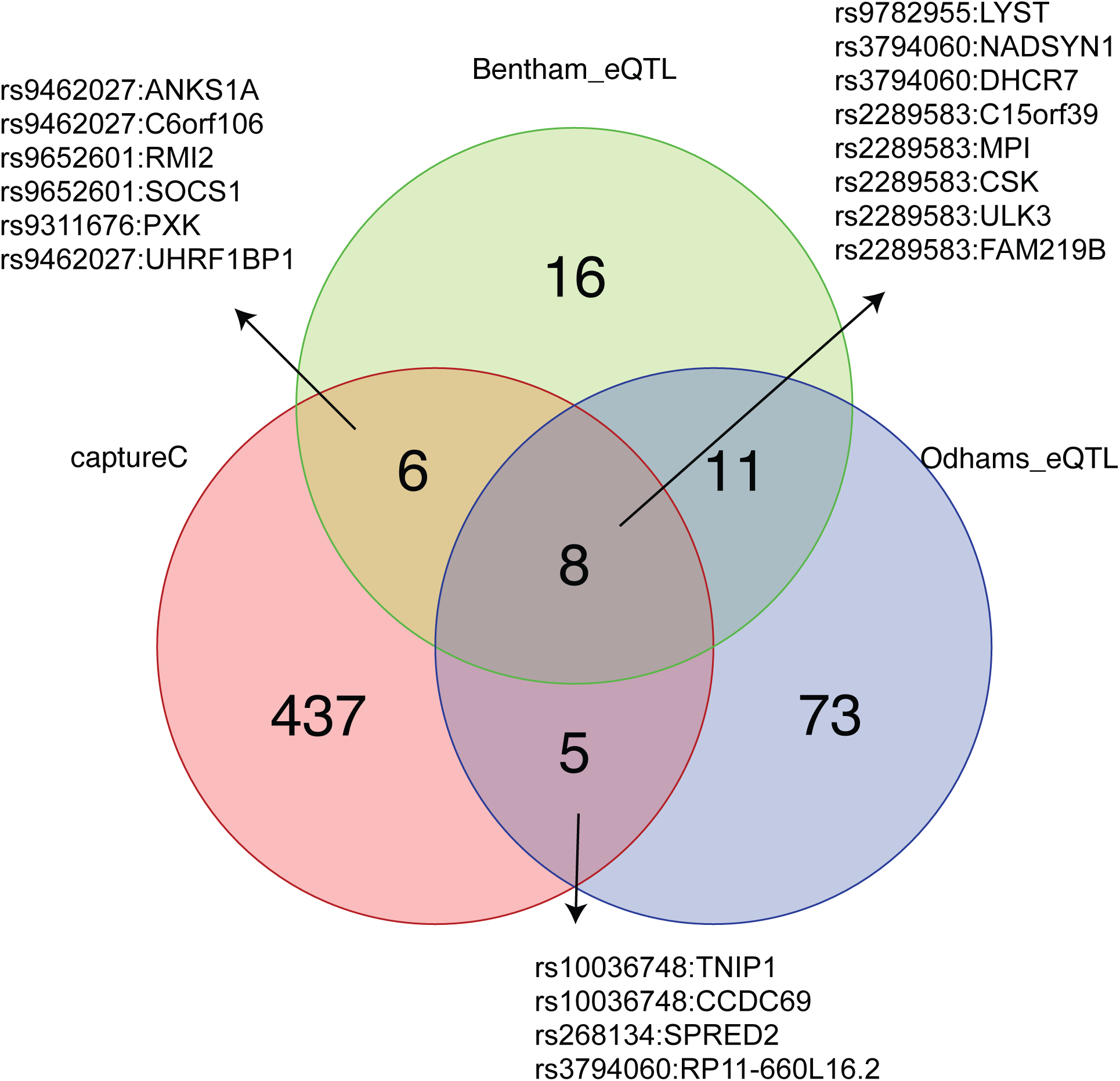
Comparison of SLE SNP-gene associations obtained by promoter-open chromatin connectomes vs. eQTL studies. Comparison of sentinel SNP-gene pairs implicated by the promoter-open chromatin connectomes in this study vs. sentinel SNP-gene pairs statistically associated in two SLE eQTL studies7,29. SNP-gene pairs shared by each group are detailed.

**SUPPLEMENTARY FIGURE 8.**
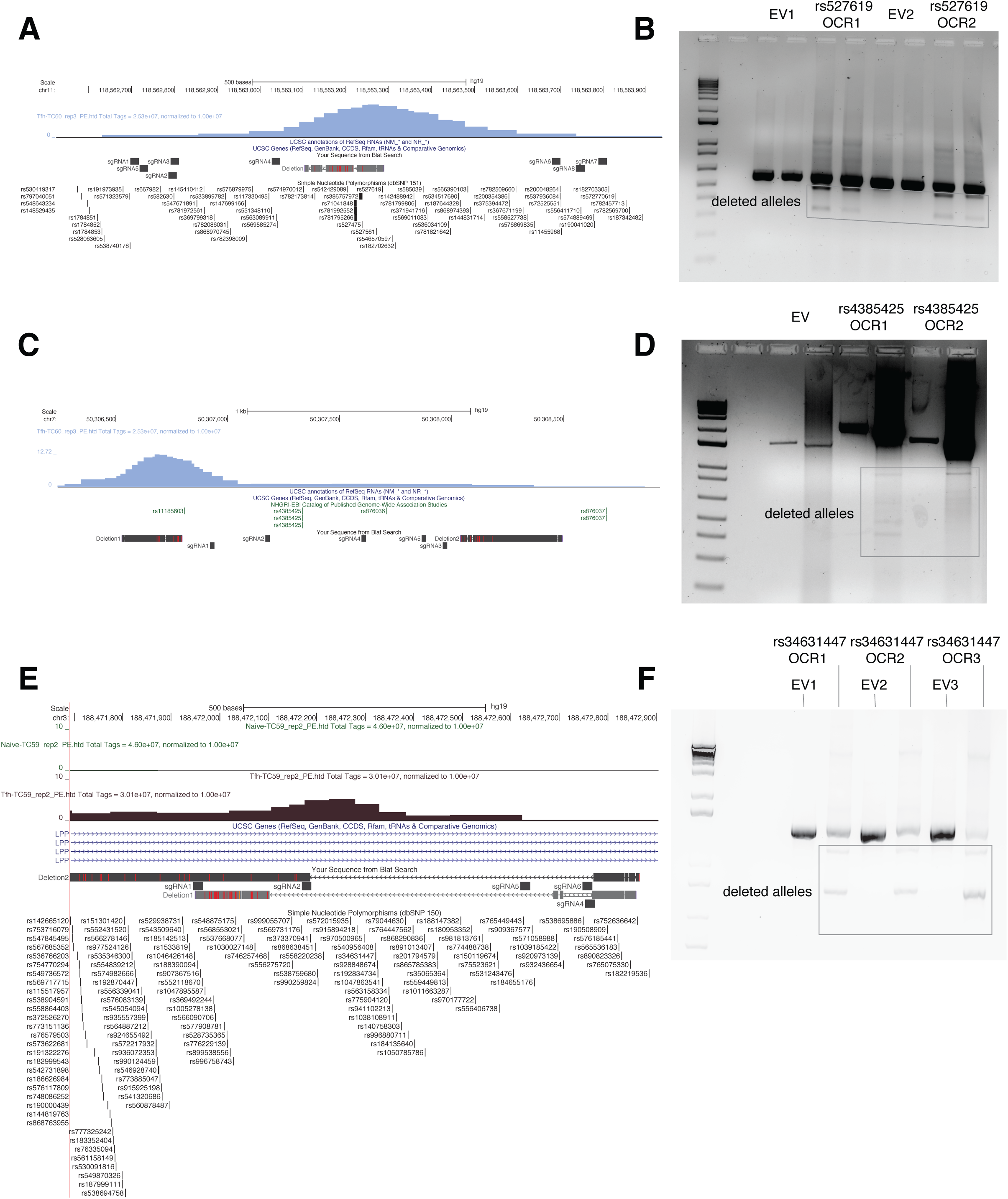
CRISPR/CAS9-based editing of accessible genomic regions containing SLE GWAS proxy SNP in Jurkat T cells. a, UCSC genome browser track displaying intergenic TFH OCR near the TREH gene harboring the rs527619 and rs71041848 proxies to the rs4639966 SLE sentinel SNP that interacts with CXCR5 (chr11:118,563,185-118,563,321). Eight sgRNAs flanking the OCR and Sanger sequencing identifying several deletions present within the OCR are depicted. b, Electrophoresis gel analysis of PCR amplified regions encompassing the targeted region showing two distinct deletions present at 500bp and 350bp. c, Genomic region surrounding the TFH OCR containing the rs4385425 SNP proxy to the sentinel SLE SNP rs11185603 (chr7:50307234-50307447) that interacts with IKZF1, showing sgRNAs and deletions detected in targeted Jurkat lines. Note that this OCR was called by HOMER, but not by MACS2. d, Electrophoresis gel analysis detects three different deletions at 900bp, 400bp and 350bp. e, Intronic OCR (chr3:188,472,234-188,472,390) in the LPP locus harboring the rs34631447 and rs79044630 SNPs proxy to sentinel rs6762714 SLE SNP and found connected to BCL6. This region was targeted with five total sgRNAs surrounding the OCR and Sanger sequencing showed two distinct deletions. f, Electrophoresis gel analysis detects 1200bp and 821bp deletions. All experiments were performed in three biological replicates.

**SUPPLEMENTARY FIGURE 9.**
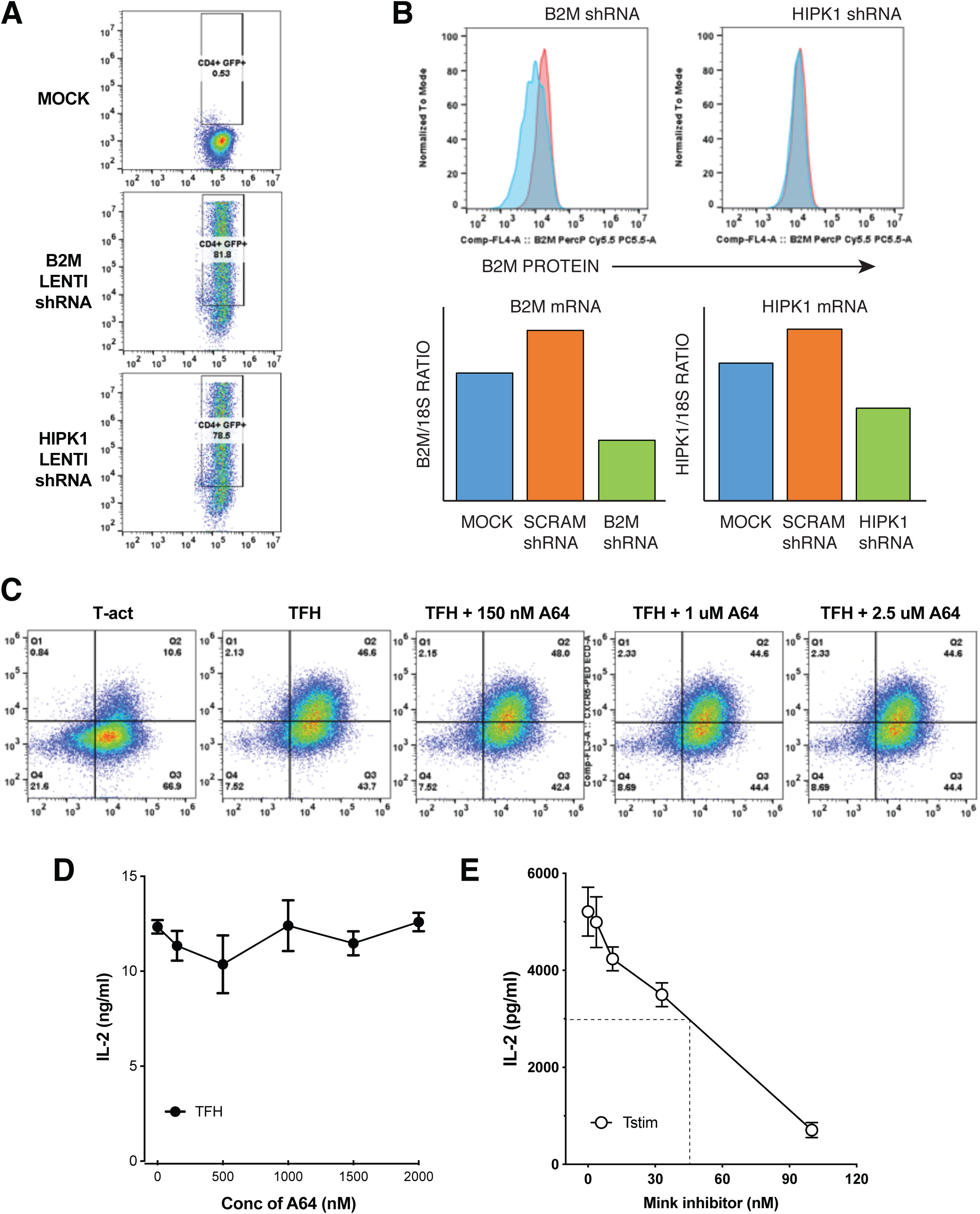
Promoter-variant connectome-guided targeting of novel kinases for modulation of primary human TFH function. a, Lentiviral delivery of B2M shRNA and HIPK1 shRNA into in vitro differentiated TFH as assessed by GFP fluorescence by flow cytometry. b, Assessment of shRNA-mediated knock-down of B2M and HIPK1 in TFH by flow cytometry and qRT-PCR. Red histograms are TFH transduced with scrambled control shRNA, and blue histograms depict TFH transduced with specific B2M (left panel) or HIPK1 (right panel) shRNA. c, Effect of HIPK inhibitory drug treatment on TFH differentiation in vitro as measured by co-induction of PD-1 and CXCR5. d, The HIPK inhibitory drug A64 does not affect IL-2 secretion by TFH cells as measured by ELISA. e, A MINK inhibitory drug inhibits IL-2 secretion by activated T cells with an ED50 of ∼50 nM. All data are representative of 3-4 replicate experiments.

**SUPPLEMENTARY FIGURE 10.**
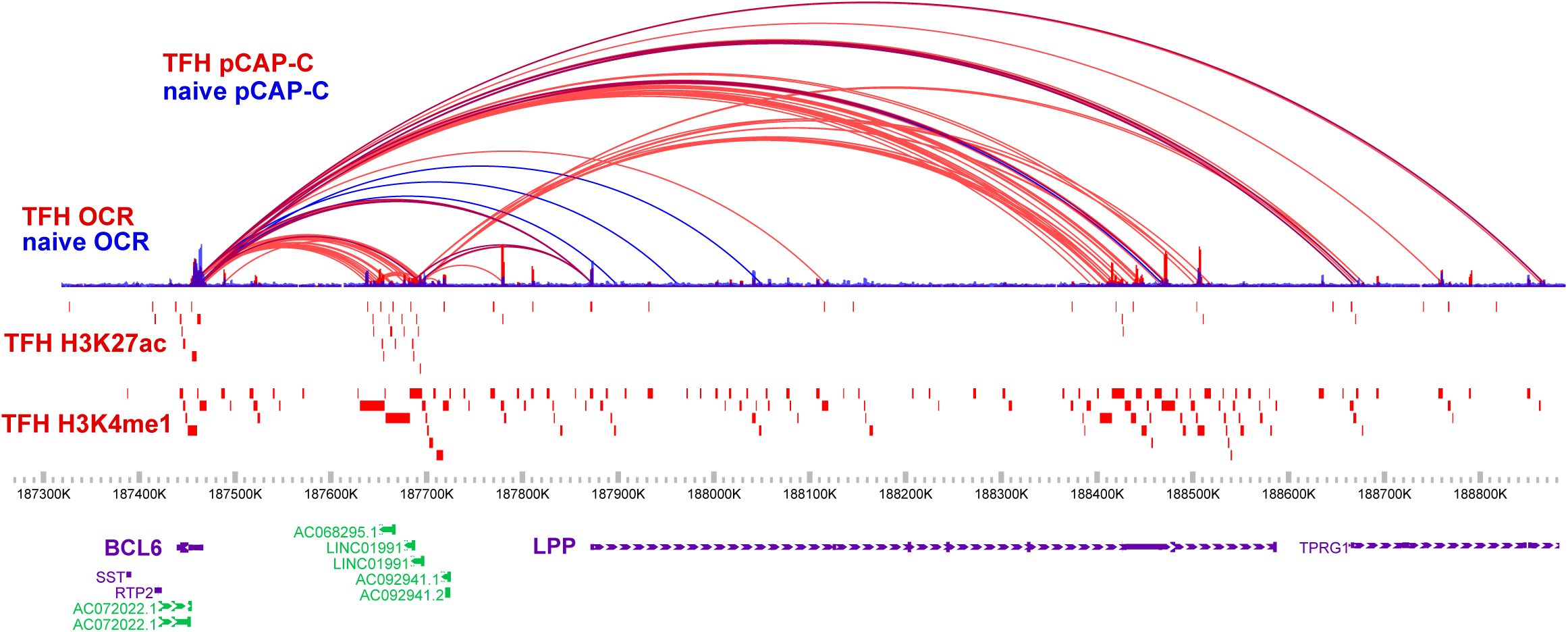
3D epigenomic map of promoter-Capture-C, ATAC-seq, H3K27ac, and H3K4me1 in the BCL6-LPP region in naïve (blue) and TFH cells (red).

## REFERENCES

1. Smemo, S. et al. Obesity-associated variants within FTO form long-range functional connections with IRX3. Nature 507, 371–375 (2014).

2. Claussnitzer, M. et al. FTO Obesity Variant Circuitry and Adipocyte Browning in Humans. New Engl J Medicine 373, 895–907 (2015).

3. Xia, Q. et al. The type 2 diabetes presumed causal variant within TCF7L2 resides in an element that controls the expression of ACSL5. Diabetologia 59, 2360–2368 (2016).

4. Tsokos, G. C., Lo, M. S., Reis, P. & Sullivan, K. E. New insights into the immunopathogenesis of systemic lupus erythematosus. Nat Rev Rheumatol 12, 716–730 (2016).

5. Song, W. & Craft, J. T follicular helper cell heterogeneity: Time, space, and function. Immunol Rev 288, 85–96 (2019).

6. Crotty, S. Follicular Helper CD4 T Cells (TFH). Annu Rev Immunol 29, 621–663 (2011).

7. Bentham, J. et al. Genetic association analyses implicate aberrant regulation of innate and adaptive immunity genes in the pathogenesis of systemic lupus erythematosus. Nat Genet 47, 1457–1464 (2015).

8. Morris, D. L. et al. Genome-wide association meta-analysis in Chinese and European individuals identifies ten new loci associated with systemic lupus erythematosus. Nat Genet 48, 940–946 (2016).

9. Thurman, R. E. et al. The accessible chromatin landscape of the human genome. Nature 489, 75 (2012).

10. Romberg, N. et al. CVID-associated TACI mutations affect autoreactive B cell selection and activation. J Clin Invest 123, 4283–4293 (2013).

11. Buenrostro, J. D., Giresi, P. G., Zaba, L. C., Chang, H. Y. & Greenleaf, W. J. Transposition of native chromatin for fast and sensitive epigenomic profiling of open chromatin, DNA-binding proteins and nucleosome position. Nat Methods 10, 1213–1218 (2013).

12. Adorini, L. & Penna, G. Control of autoimmune diseases by the vitamin D endocrine system. Nat Clin Pract Rheum 4, 404 EP–412 (2008).

13. Hughes, J. R. et al. Analysis of hundreds of cis-regulatory landscapes at high resolution in a single, high-throughput experiment. Nat Genet 46, 205–212 (2014).

14. Chesi, A. et al. Genome-scale Capture C promoter interactions implicate effector genes at GWAS loci for bone mineral density. Nat Commun 10, 1260 (2019).

15. Wingett, S. et al. HiCUP: pipeline for mapping and processing Hi-C data. F1000research 4, 1310 (2015).

16. Cairns, J. et al. CHiCAGO: robust detection of DNA looping interactions in Capture Hi-C data. Genome Biol 17, 127 (2016).

17. Javierre, B. M. et al. Lineage-Specific Genome Architecture Links Enhancers and Non-coding Disease Variants to Target Gene Promoters. Cell 167, 1369–1384.e19 (2016).

18. Ernst, J. et al. Mapping and analysis of chromatin state dynamics in nine human cell types. Nature 473, 43–49 (2011).

19. Weinstein, J. S. et al. Global transcriptome analysis and enhancer landscape of human primary T follicular helper and T effector lymphocytes. Blood 124, 3719–3729 (2014).

20. Bernstein, B. E. et al. The NIH Roadmap Epigenomics Mapping Consortium. Nat Biotechnol 28, 1045 (2010).

21. Cuddapah, S. et al. Global analysis of the insulator binding protein CTCF in chromatin barrier regions reveals demarcation of active and repressive domains. Genome Res 19, 24–32 (2009).

22. Johnston, R. J. et al. Bcl6 and Blimp-1 Are Reciprocal and Antagonistic Regulators of T Follicular Helper Cell Differentiation. Science 325, 1006–1010 (2009).

23. Kroenke, M. A. et al. Bcl6 and Maf Cooperate To Instruct Human Follicular Helper CD4 T Cell Differentiation. J Immunol 188, 3734–3744 (2012).

24. Hatzi, K. et al. BCL6 orchestrates Tfh cell differentiation via multiple distinct mechanisms. J Exp Medicine 212, 539–553 (2015).

25. Liu, X. et al. Genome-wide Analysis Identifies Bcl6-Controlled Regulatory Networks during T Follicular Helper Cell Differentiation. Cell Reports 14, 1735–1747 (2016).

26. Liu, X. et al. Bcl6 expression specifies the T follicular helper cell program in vivo. J Exp Medicine 209, 1841–1852 (2012).

27. Eystathioy, T., Peebles, C. L., Hamel, J. C., Vaughn, J. H. & Chan, E. K. Autoantibody to hLSm4 and the heptameric LSm complex in anti-Sm sera. Arthritis Rheumatism 46, 726–734 (2002).

28. Gunnewiek, K. J., van de Putte, L. & van Venrooij, W. The U1 snRNP complex: an autoantigen in connective tissue diseases. An update. Clin Exp Rheumatol 15, 549–60 (1997).

29. Odhams, C. A. et al. Mapping eQTLs with RNA-seq reveals novel susceptibility genes, non-coding RNAs and alternative-splicing events in systemic lupus erythematosus. Hum Mol Genet 26, 1003–1017 (2017).

30. Kim, Y., Choi, C., Lee, S., Conti, M. & Kim, Y. Homeodomain-interacting protein kinases, a novel family of co-repressors for homeodomain transcription factors. J Biol Chem 273, 25875–25879 (1998).

31. Nicke, B. et al. Involvement of MINK, a Ste20 Family Kinase, in Ras Oncogene-Induced Growth Arrest in Human Ovarian Surface Epithelial Cells. Mol Cell 20, 673–685 (2005).

32. Locci, M. et al. Activin A programs the differentiation of human TFH cells. Nat Immunol 17, 976–984 (2016).

33. Rubin, A. J. et al. Lineage-specific dynamic and pre-established enhancer-promoter contacts cooperate in terminal differentiation. Nat Genet 49, (2017).

34. o, L. T. et al. Genome-wide characterization of mammalian promoters with distal enhancer functions. Nat Genet 49, 1073–1081 (2017).

35. Sanyal, A., Lajoie, B. R., Jain, G. & Dekker, J. The long-range interaction landscape of gene promoters. Nature 489, 109 (2012).

36. Bunting, K. L. et al. Multi-tiered Reorganization of the Genome during B Cell Affinity Maturation Anchored by a Germinal Center-Specific Locus Control Region. Immunity 45, 497–512 (2016).

37. Kondo, S. et al. Characterization of cells and gene-targeted mice deficient for the p53-binding kinase homeodomain-interacting protein kinase 1 (HIPK1). Proc National Acad Sci 100, 5431–5436 (2003).

38. Louie, S. H. et al. Modulation of the beta-catenin signaling pathway by the dishevelled-associated protein Hipk1. Plos One 4, e4310 (2009).

39. Shang, Y. et al. Transcriptional corepressors HIPK1 and HIPK2 control angiogenesis via TGF-β-TAK1-dependent mechanism. Plos Biol 11, e1001527 (2013).

40. Guerra, F. M., Gommerman, J. L., Corfe, S. A., Paige, C. J. & Rottapel, R. Homeodomain-Interacting Protein Kinase (HIPK)-1 Is Required for Splenic B Cell Homeostasis and Optimal T-Independent Type 2 Humoral Response. Plos One 7, e35533 (2012).

41. Larhammar, M., Huntwork-Rodriguez, S., Rudhard, Y., Sengupta-Ghosh, A. & Lewcock, J. W. The Ste20 Family Kinases MAP4K4, MINK1, and TNIK Converge to Regulate Stress-Induced JNK Signaling in Neurons. J Neurosci 37, 11074–11084 (2017).

42. Kaneko, S. et al. Smad inhibition by the Ste20 kinase Misshapen. Proc National Acad Sci 108, 11127–11132 (2011).

43. Fu, G. et al. Suppression of Th17 cell differentiation by misshapen/NIK-related kinase MINK1. J Exp Med 214, jem.20161120 (2017).

44. Ozaki, K. et al. A Critical Role for IL-21 in Regulating Immunoglobulin Production. Science 298, 1630–1634 (2002).

45. Sheffield, N. C. & Bock, C. LOLA: enrichment analysis for genomic region sets and regulatory elements in R and Bioconductor. Bioinformatics 32, 587–589 (2016).

